# Hidden Dynamical Canalization at the Onset of *Hydra* Morphogenesis

**DOI:** 10.64898/2026.04.28.721438

**Authors:** Oded Agam, Erez Braun

## Abstract

The primary morphological transition in *Hydra* regeneration, from a quasi-spherical fragment to an elongated body, is preceded by a prolonged period during which its average form changes only modestly, while its instantaneous shape continues to fluctuate. We asked whether this preparatory stage contains signatures of dynamical organization. Using principal component analysis, we calculated the effective dimension of shape fluctuations, which characterizes how broadly they are distributed across PCA directions. Across all five conditions, spanning differences in tissue fragment size, axial origin and GdCl₃ exposure, the effective dimension decreased progressively before overt tissue elongation. Total fluctuation variance increased about 2.8-fold, whereas the leading eigenvalue increased about fourfold. Late intervals had lower effective dimension than early intervals at similar coarse shapes. Calcium analysis in GdCl₃-treated tissues distinguished mean level, fast events and spatial heterogeneity. Fast events were associated with short-timescale shape changes. After accounting for temporal trends, mean Ca^2+^ and coarse morphology, higher spatial heterogeneity of Ca^2+^ activity was associated with lower effective dimension. The progressive concentration of morphological fluctuations along fewer dominant directions characterizes dynamical canalization associated with spatial Ca^2+^ heterogeneity.

**SUMMARY STATEMENT:** Before visible elongation, regenerating *Hydra* tissues progressively concentrate their shape fluctuations along fewer dominant directions, revealing dynamical organization during the early quasi-spherical period.

## INTRODUCTION

Animal morphogenesis is the process by which a complex stereotypical body form emerges from an initially simpler state. This process is remarkable for its robustness, producing reliable functional structures under fluctuating, heterogeneous and noisy conditions^1^. Waddington coined the term canalization to describe this robustness, proposing that it arises from system dynamics directed by a static landscape of steep canals corresponding to developmental pathways, shaped by epigenetic processes^2,3^. While this concept has been highly successful in explaining canalization in cell differentiation and tissue patterning, the generation of body form also requires mechanical processes that organize morphology^4,5^. In many systems, the visible geometry of the body reflects only part of the developmental process, which integrates biochemical, mechanical and electrical activities tightly coupled across scales^5–15^. This raises the possibility that morphogenetic organization may be expressed by morphological fluctuations before overt shape changes. Here, we use the concept of dynamical canalization in a narrow operational sense to describe progressive concentration of shape-fluctuation variance into fewer dominant modes, accompanied by temporal organization of the leading mode. This concept is distinct from the classical Waddingtonian sense of robustness of the developmental outcome. Utilizing whole-body *Hydra* regeneration from a tissue fragment, we ask whether such organization emerges during the early quasi-spherical period preceding the primary morphological transition to an elongated tube-like structure, the hallmark of a mature *Hydra*.

*Hydra* regeneration from a tissue fragment provides a particularly useful system for addressing the issue of morphological canalization^5,7,14,16–20^. Following excision, a tissue fragment folds into a closed hollow spheroid and later develops an elongated morphology along a defined body axis. Regeneration proceeds largely without substantial cell proliferation^21–23^ and is strongly influenced by active actomyosin-generated forces^7,20^ and hydrostatic pressure, modulated by osmotic pressure gradients, within the enclosed lumen^24–26^. Several processes take place during the early stages of regeneration. The epithelial supracellular actin fibers become organized into an ordered array^7,27^, transient contractions focus at the actin defect site, and the head organizer emerges at this defect location^20,28^. At the same time, water inflow driven by the osmotic pressure gradient increases lumen pressure, leading to occasional tissue rupture-repair cycles manifested in slow increase followed by sharp drops in the tissu’e projected area^24,28,29^. Importantly, tissue fragments excised under different initial and physiological conditions can all regenerate into mature *Hydra*, even though their morphogenetic trajectories are not identical^16^. For example, we previously showed that a small fragment excised from the central gastric region (mid-axis) often remains nearly spherical for an extended interval, which we termed the “preparatory period”, before undergoing a relatively rapid transition to an elongated, tube-like morphology^13,14^. In contrast, large central fragments, as well as small fragments excised near the head or foot, pass through a similar early quasi-spherical preparatory period but then reshape more gradually into an elongated form, with broader fluctuations and transient intermediate morphologies^16^. Thus, tissues that appear similar in overall shape during the early preparatory period may nevertheless differ in their inherited positional information and in the subsequent route by which regeneration proceeds.

While our previous work provided a comprehensive experimental and theoretical framework for explaining the major morphological transition under different initial and physiological conditions, the preparatory period remains poorly characterized^13,14,16,30^. Here, we focus on this period, looking for hidden dynamical changes that precede the main transition to a tube-like shape. During this interval, the tissue fluctuates around a quasi-spherical morphology, and coarse shape descriptors show only modest changes. Yet, weak changes in coarse morphology do not necessarily imply that the underlying dynamical state remains unchanged. The central question, therefore, is whether this early quasi-spherical regime is simply a passive fluctuating state, or whether it already reflects the onset of dynamical canalization that precedes the major morphological transition.

To address this question, we analyze the structure of shape fluctuations during the preparatory time window using principal component analysis (PCA). We use the effective dimension, derived from the spectral entropy of the normalized PCA variance spectrum, to characterize how broadly shape fluctuations are distributed across PCA directions ^31^. This framework allows us to ask whether tissues with similar coarse morphology can differ in their fluctuation spectra and how these spectra vary with positional cues inherited from the donor animal and with physiological perturbation. Building on our previous work demonstrating the central role of Ca²⁺ in the morphological transition^10,11,13,14,16^, we examine how spatial features of calcium activity relate to the effective dimension of shape fluctuations during the preparatory period.

Here, we show that the quasi-spherical preparatory stage is not dynamically stationary. PCA identifies a progressive redistribution of shape fluctuations toward fewer dominant modes, which we interpret as an operational signature of dynamical canalization before overt elongation. Comparisons among fragments differing in size, axial origin and GdCl₃ exposure that blocks Ca^2+^ channels, reveal condition-dependent differences in this organization. In Ca^2+^ recordings from GdCl₃-treated tissues, higher spatial heterogeneity of calcium activity was associated with a lower effective dimension. Because GdCl_3_ is widely used as a blocker of stretch-activated mechanosensitive ion channels and can also inhibit certain other Ca^2+^ channels, its action is expected to reduce mechanically induced calcium response. Calcium signals, in turn, are known to promote actomyosin activity and contractile force generation^10,32–35^. Indeed, we previously showed that GdCl_3_ treated tissue samples exhibit slow-down responses to tissue ruptures affecting their repair dynamics^29^. Together, these findings indicate that fluctuation spectra can reveal morphogenetic organization that remains hidden when only coarse morphology is inspected.

## RESULTS

### The preparatory stage of morphogenesis

We analyzed time-lapse measurements of tissue fragments differing in size, axial origin and GdCl_3_ exposure. The data from the untreated conditions were obtained in previously reported experiments, whereas the data for the nine GdCl_3_-treated tissues are new to the present study. The comparative analysis of contour fluctuations during the early quasi-spherical period is new for all samples under the five conditions. Dataset provenance and characteristics are summarized in Table S1.

The preparatory stage was identified using the shape parameter previously employed to characterize the tissue morphology, *Λ = 1 − 4πA/P^2^*, where *A* is the projected area of the tissue and *P* its perimeter^13,14,16^. This parameter is zero for a circular projected image and increases with any departure from circularity, including elongation and irregular contour deformations. It provides a broad scalar measure sensitive to multiple modes of projected morphological deformation. We, therefore, operationally define the preparatory stage as the time interval during which the shape parameter remains low, typically Λ(t)≲0.15, before the onset of the primary morphological transition. This criterion is utilized as a practical guide rather than a strict cutoff: brief excursions above this value may occur, while the tissue still remains within the preparatory regime. Our conclusions are not sensitive to the exact choice of this threshold. The complete shape-parameter trajectories of the nine GdCl₃-treated tissues, together with the tissue-specific endpoints of the analyzed early quasi-spherical period, are shown in Fig. S1. Corresponding trajectories for the remaining tissue datasets analyzed here were reported previously in the Supplementary Information of Refs.^13,14,16^.

Figure 1A shows a representative time trace of the shape parameter for a small GdCl_3_-treated tissue fragment. The black segment marks the preparatory stage, during which Λ(t) remains low, whereas the red segment marks the interval in which the main morphological transition occurs. The corresponding projected contours, shown in Fig. 1B, illustrate the tissue’s morphology during this low-Λ regime: throughout the preparatory stage, the tissue fluctuates continuously while remaining centered around a broadly quasi-spherical shape. Only later, during the interval corresponding to the red segment in Fig. 1A, does the tissue undergo a pronounced elongation, which is clearly visible in the projected contours.

**Fig. 1.**
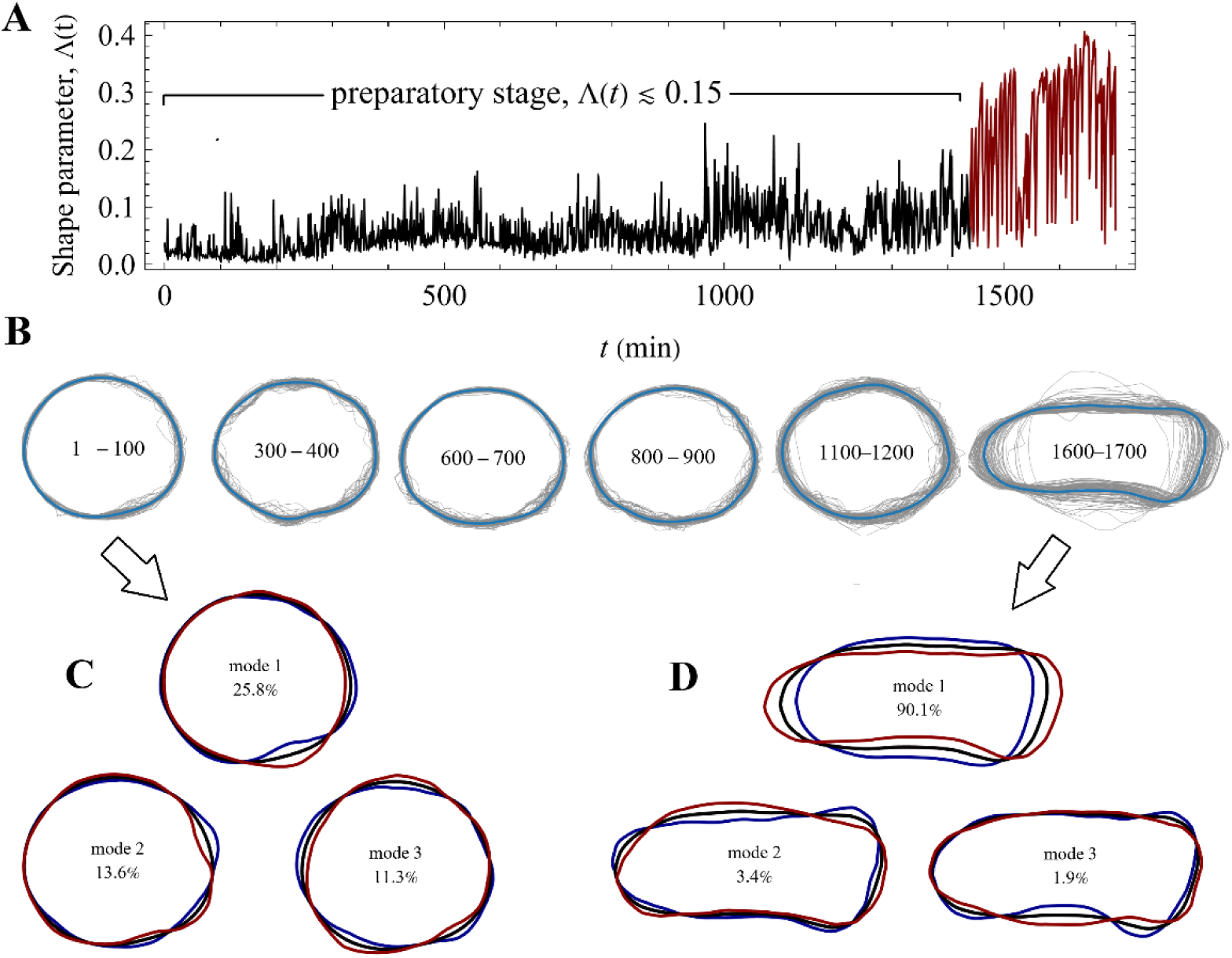
The preparatory stage in a representative GdCl_3_-treated small tissue fragment. **(A)** Time evolution of the shape parameter Λ(t) for a representative tissue. Low Λ indicates a nearly circular projected contour. The black solid line marks the preparatory stage, during which Λ(t) remains low, whereas the dark red line marks the interval in which the primary morphological transition occurs. **(B)** Representative area-normalized and aligned projected contours in successive time windows (see Methods). Gray lines show individual contours, and the blue line shows the local average shape within each window. During the preparatory stage the tissue fluctuates continuously around a broadly quasi-spherical morphology. Pronounced elongation appears only later, during the interval corresponding to the red segment in panel A. **(C)** PCA modes computed from an early preparatory window using the same preprocessed contour representation (see Methods). The black curve denotes the mean shape in the selected window, and the red and blue curves indicate opposite deformations along each PCA mode. Here, the variance is distributed over several modes, with the first three modes accounting for 25.8%, 13.6%, and 11.3% of the variance, respectively. **(D)** PCA modes computed from a late post-transition window, using the same preprocessed contour representation. The black curve again denotes the mean shape, and the red and blue curves indicate opposite deformations along each PCA mode. After the morphological transition, the fluctuations are strongly concentrated in the first mode, which accounts for 90.1% of the variance, whereas the relative contributions of higher modes are strongly reduced, consistent with a marked decrease in the effective dimension of the morphological fluctuations.

To characterize the structure of these fluctuations, we applied PCA to preprocessed projected contours within selected time windows (see Methods). Before PCA, contours were rescaled to unit area, translated to a common centroid, aligned to a common orientation, and cyclically reindexed to remove phase differences along the contour (see Methods). This preprocessing ensured that the PCA captures shape fluctuations rather than trivial variation due to inflation, position, orientation, or contour parametrization. During the early preparatory stage, the fluctuations are distributed over several modes of comparable weights. In the representative case of Fig. 1A, the first three modes, shown in panel C, account for 25.8%, 13.6%, and 11.3% of the variance, respectively. By contrast, after the morphological transition, the fluctuations become strongly concentrated in the first mode, which alone accounts for 90.1% of the variance, while the fractions of total variance carried by the higher modes are strongly reduced. To represent this redistribution of variances, we denote by *λ_k_* the eigenvalue associated with PCA mode *k*, representing the variance carried by the direction of the corresponding eigenvector, and define *p_k_* = *λ_k_* _∑_*_j_ λ_j_* as its fraction of the total variance in projected shape fluctuations. The spectral entropy *H* = −_∑_*_k_ p_k_* ln*p_k_* defines an effective dimension *d*_eff_ = exp(*H*), which quantifies the number of PCA modes carrying appreciable variance in the scale-normalized contour representation^31^. Specifically, the effective dimension associated with the first tissue contour set shown in panel B is 8.16, whereas that of the last contour set is only 1.23, consistent with a strong restriction of the observed fluctuations to essentially a single dominant PCA mode after the morphological transition.

While the reduction in effective dimension between the preparatory stage and the post-transition regime is not surprising, it is unclear whether a similar reduction also occurs within the preparatory stage itself, where the tissue remains essentially quasi-spherical. If so, this would indicate that the approach to the morphological transition already involves an internal morphological dynamical reorganization before pronounced shape change becomes visible. The preparatory stage would then correspond not to a stationary fluctuation regime around an almost unchanged morphology, but to a period of progressive dynamical canalization within an apparently similar coarse shape.

### Effective dimension declines within the preparatory stage

To test whether dynamical canalization is already underway during the preparatory stage, we examined how the effective dimension evolves within this interval. Because regeneration trajectories vary substantially, even among tissue samples of the same type or condition, comparing samples at matched absolute times would not align with equivalent dynamical phases^16^. Figure 2A summarizes the procedure: the preparatory-stage duration of each tissue was normalized, divided into 10 equal intervals, and analyzed by PCA independently within each interval to calculate *d*_eff_ from the corresponding eigenspectrum (see Methods). This analysis reveals a consistent decline in *d*_eff_ as the preparatory stage progresses.

**Fig. 2.**
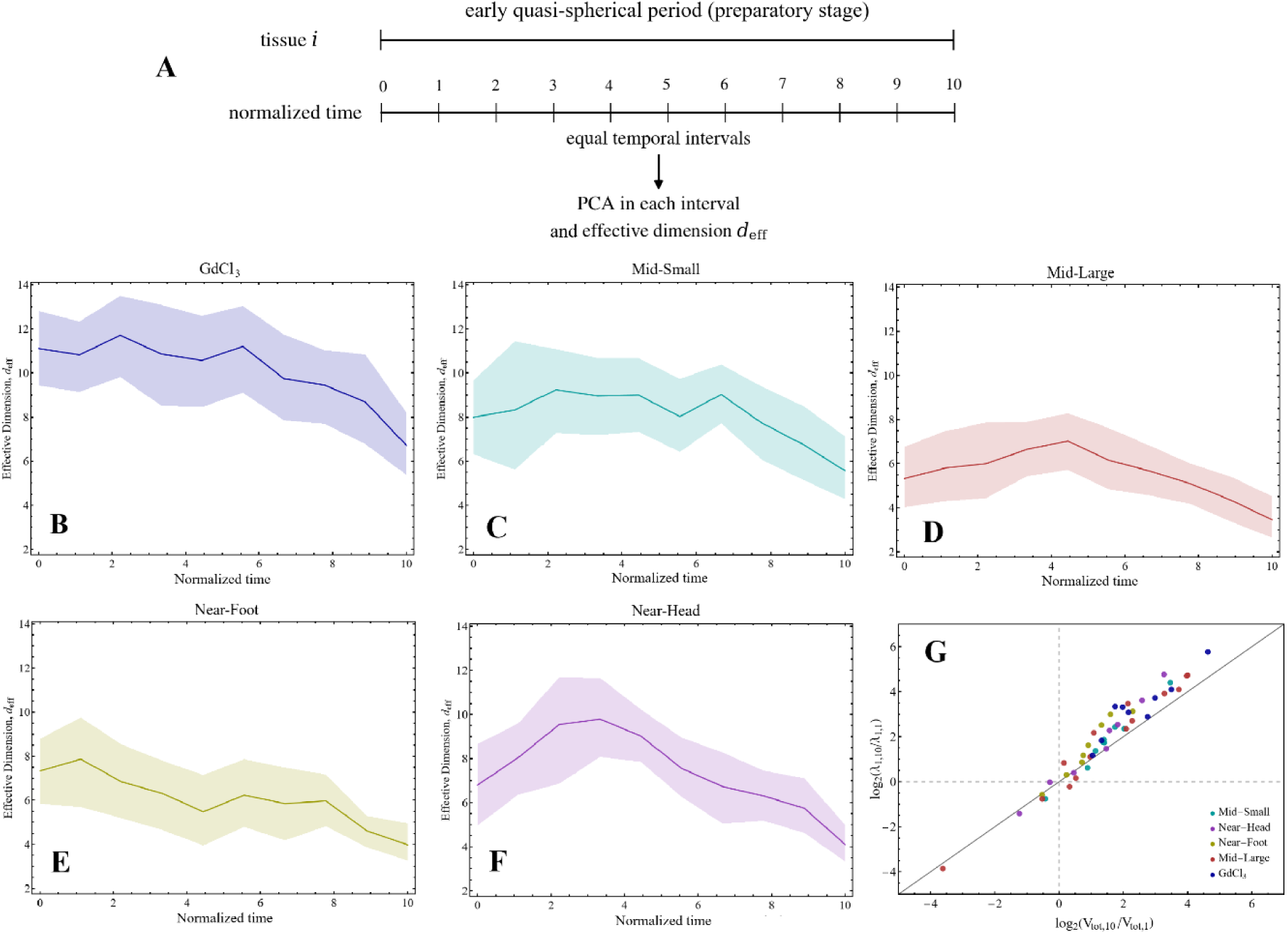
Shape-fluctuation variance becomes progressively concentrated during the early quasi-spherical period. (A) Schematic of the analysis. For each tissue, the early quasi-spherical period was normalized in time and divided into 10 equal intervals. PCA was performed independently within each interval, and the effective dimension, *d*_eff_, was calculated from the corresponding eigenspectrum. Lower *d*_eff_ indicates that variance of the projected shape fluctuations is concentrated into fewer PCA directions. (B–F) Mean *d*_eff_ as a function of normalized time for GdCl₃, Mid-Small, Mid-Large, Near-Foot and Near-Head tissues, respectively. Solid lines show means across tissues and shaded regions show tissue-bootstrap 95% confidence intervals. (G) Animal-level comparison of the initial-to-final fold changes in the leading PCA eigenvalue, *λ*_1_, and total variance, *V*_tot_ = _∑_*_k_ λ_k_*. Each symbol represents one tissue sample and colors identify the five experimental conditions. Both axes show the base-2 logarithm of the ratio between the final and initial intervals. The diagonal denotes equal proportional changes in *λ*_1_ and V_tot_; points above it indicate disproportionate amplification of the variance along the leading fluctuation direction.

Averaging over samples within each condition shows that this decline is a general feature of the preparatory stage rather than a property of a single tissue class (Fig. 2B-F). In every condition, *d*_eff_ is higher at early times and lower near the end of the preparatory window, indicating that fluctuation variance becomes progressively concentrated into fewer PCA directions before the morphological transition. Thus, even while the tissue remains approximately quasi-spherical, the fluctuation dynamics are not stationary. Instead, they become increasingly confined to a smaller set of collective modes as the transition is approached.

At the same time, the absolute level of the effective dimension depends strongly on the tissue’s initial conditions. Tissues inheriting positional cues from near the head or foot of the parent *Hydra* generally exhibit lower values of *d*_eff_, consistent with the idea that positional information constrains the fluctuation space explored by the tissue. This tendency is most evident in Mid-Large and Near-Foot tissues, which remain at comparatively low dimension values throughout most of the preparatory period, and is also reflected in the lower late-time effective dimensions reached by all unperturbed tissue classes. In this sense, positional cues appear not only to shape the large-scale morphological dynamics, but also to promote the concentration of shape fluctuations into fewer dominant directions during the preparatory stage.

By contrast, GdCl₃-treated tissues, although showing an overall decline, maintain substantially higher effective dimension throughout much of the preparatory stage (Fig. 2B). Table 1 summarizes the initial-to-final changes and the corresponding animal-level statistical analysis. Importantly, however, the magnitude and temporal profile of the decrease are dependent on the tissue’s condition.

**Table 1.** Effective dimension in the initial and final intervals of the early quasi-spherical period.

| Condition | n | Initial mean $d_{\text{eff}}$ | Final mean $d_{\text{eff}}$ | Mean paired change, $\Delta d_{\text{eff}}$ |
| --- | --- | --- | --- | --- |
| GdCl<sub>3</sub> | 9 | 11.11 | 6.73 | 4.39 |
| Mid-Small | 9 | 7.99 | 5.57 | 2.43 |
| Mid-Large | 14 | 5.32 | 3.45 | 1.86 |
| Near-Foot | 8 | 7.33 | 3.98 | 3.35 |
| Near-Head | 8 | 6.79 | 4.10 | 2.69 |

Values are means across tissue samples. $\Delta d_{\text{eff}} = d_{\text{eff}}^{\text{start}} - d_{\text{eff}}^{\text{end}}$ ; positive values indicate a decrease. Effective dimension decreased in 40 of 48 tissues. From the first to the final interval, $d_{\text{eff}}$ decreased by an average of 40.5% (condition-balanced geometric mean final-to-initial ratio, 0.595; 95% CI, 0.516–0.685; two-sided sign-flip $p < 0.0001$ ). Each condition was given equal weight in the overall estimate.

Because rupture-repair events are accompanied by rapid drops in the projected tissue area, we tested whether the associated changes could account for the decline in *d*_eff_. Contour changes between consecutive rupture events exceeded changes across the rupture-associated events in 44 of the 48 sample tissues. Moreover, the decline in *d*_eff_ remained evident in all five conditions when the analysis was repeated using intervals bounded by successive rupture events with the drop frames excluded (Figs. S2 and S3, and Supplementary Note 1). Thus, the observed spectral concentration is not an artifact of the rupture timing.

Similar trends are obtained when fluctuation dimensionality is quantified by the participation ratio of the PCA spectrum rather than by the entropy-based effective dimension (Supplementary Fig. S4). The decline accompanies an approximately 2.8-fold increase in total variance and a 4-fold increase in *λ*_1_ (Fig. 2G; Fig. S5). It is robust to changes in angular resolution and contour-centering convention and remains evident after temporal subsampling at intervals of up to 20 min (Fig. S6). The leading PCA direction also shows modest temporal stabilization across intervals (Fig. S7; Supplementary Note 2).

Taken together, these results show that the preparatory stage is not a passive waiting period preceding visible shape change. Rather, it is a phase of progressive dynamical reorganization, affected by positional cues and GdCl₃ treatment.

### The relation between the shape parameter and the effective dimension is heterogeneous

A natural question is whether the decline in effective dimension during the preparatory stage simply reflects gradual changes in coarse morphology. We therefore examined the relation between the shape parameter Λ and *d*_eff_ separately for each condition (Fig. 3). Panels A–E show the pooled interval-level data together with descriptive Pearson *r*, Spearman ρ and least-squares slopes. The pooled associations varied in sign and were generally weak.

**Fig. 3.**
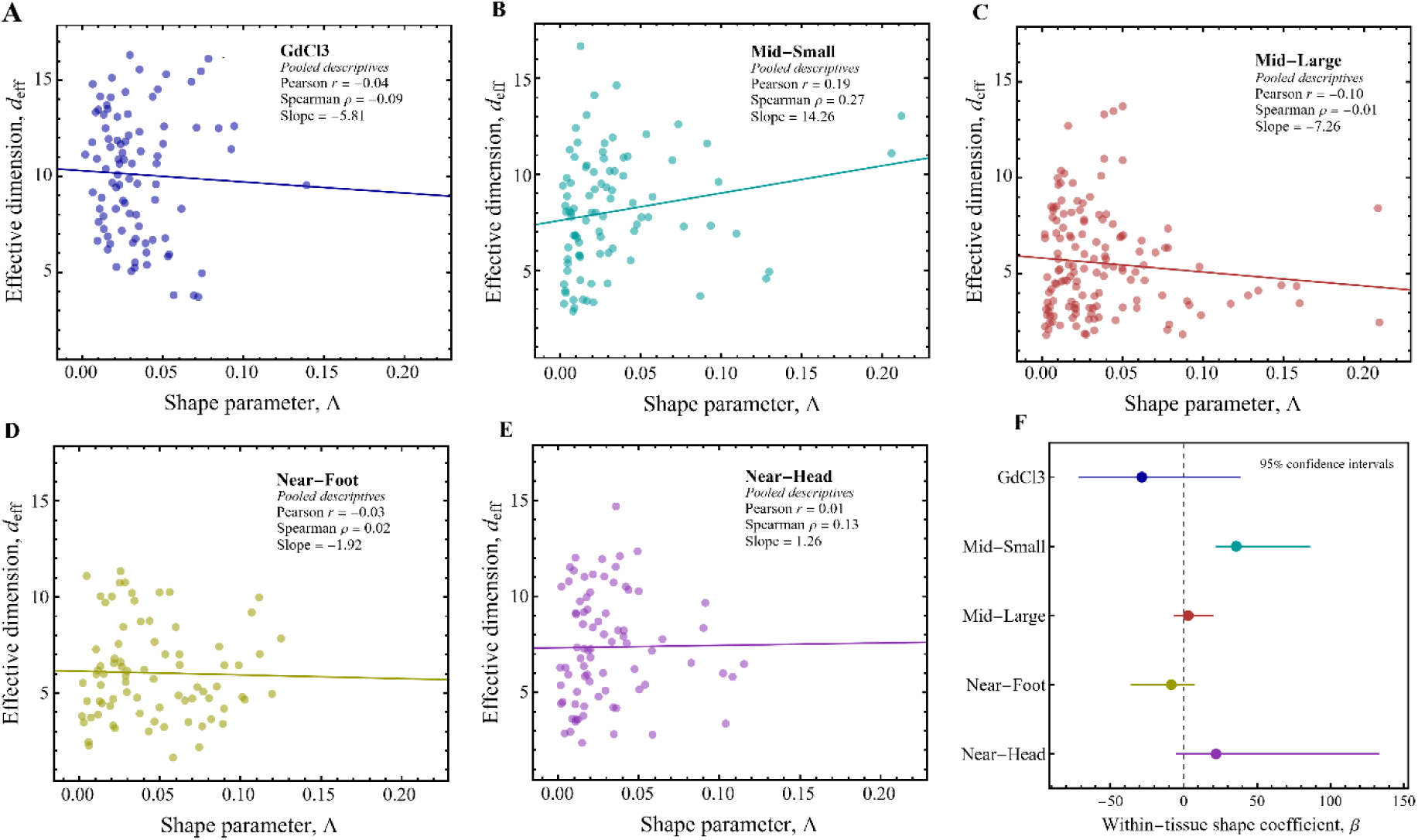
Heterogeneous relation between the shape parameter and the effective dimension. (A–E) Effective dimension, *d*_eff_, plotted against the shape parameter, Λ, separately for each condition. Each point represents one interval from one tissue sample; solid lines show pooled least-squares fits. Insets report the descriptive pooled Pearson *r*, Spearman ρ and regression slope. Because these pooled estimates combine within- and between-tissue variation, they can differ from the within-tissue estimates; statistical inference is therefore based on panel F. **(F)** Within-tissue slope, β, relating *d*_eff_ to Λ, estimated by tissue fixed-effects regression. Points show coefficient estimates, and horizontal lines show 95% confidence intervals obtained by resampling complete tissues. Exact two-sided tissue-level sign-flip P values were 0.367, 0.035, 0.568, 0.344 and 0.258 for GdCl₃, Mid-Small, Mid-Large, Near-Foot and Near-Head, respectively. None remained significant after Holm correction.

To isolate the within-tissue association, we used the tissue fixed-effects model *d*_eff_ = α_tissue_ + βΛ, where α_tissue_ accounts for tissue-specific baselines and β measures the association between Λ and *d*_eff_ within tissues (Fig. 3F). Confidence intervals were obtained by bootstrap resampling of complete tissues, and P values by exact two-sided tissue-level sign-flip tests. Pooled and within-tissue estimates differed in some conditions. For example, the negative pooled slope in Mid-Large was not reproduced within tissues, demonstrating that pooled trends can be influenced by between-tissue variation.

Only the Mid-Small condition showed evidence of a within-tissue association before multiple-testing correction (sign-flip P = 0.035); none of the five coefficients remained significant after Holm correction. Thus, Λ does not consistently predict *d*_eff_ within tissues, and the decline in effective dimension cannot be reduced to generic tracking of coarse shape.

### Similar coarse morphologies can correspond to different dynamical states

Although Λ does not uniquely determine *d*_eff_, its overall decline could still be driven mainly by a relatively small number of intervals with stronger shape changes, rather than by dynamical reorganization within broadly similar morphologies. We therefore asked whether similar coarse shapes can occupy different dynamical states during the early quasi-spherical period, consistent with the concept of dynamical canalization.

To address this question, we constructed maps of *d*_eff_ in a low-dimensional space of projected shapes. The local mean contours are approximated by a two-parameter family defined by *e*, which captures the leading anisotropy, and *s*, which captures a symmetric shoulder-like deviation from an ellipse (Fig. 4A; see Methods). Points in this space were obtained from windows of approximately 100 consecutive frames and associate a mean contour with the effective dimension of the fluctuations around it. Maps for GdCl₃-treated tissues at the beginning and end of the preparatory period are shown in Fig. 4B,C, with the corresponding shape matching in Fig. 4D. Maps and matches for the other conditions are shown in Fig. S8, while Fig. S9 presents the mean contours of an independent subset of matched pairs. Full statistical results and sensitivity to the matching tolerance are given in Table S2.

**Fig. 4.**
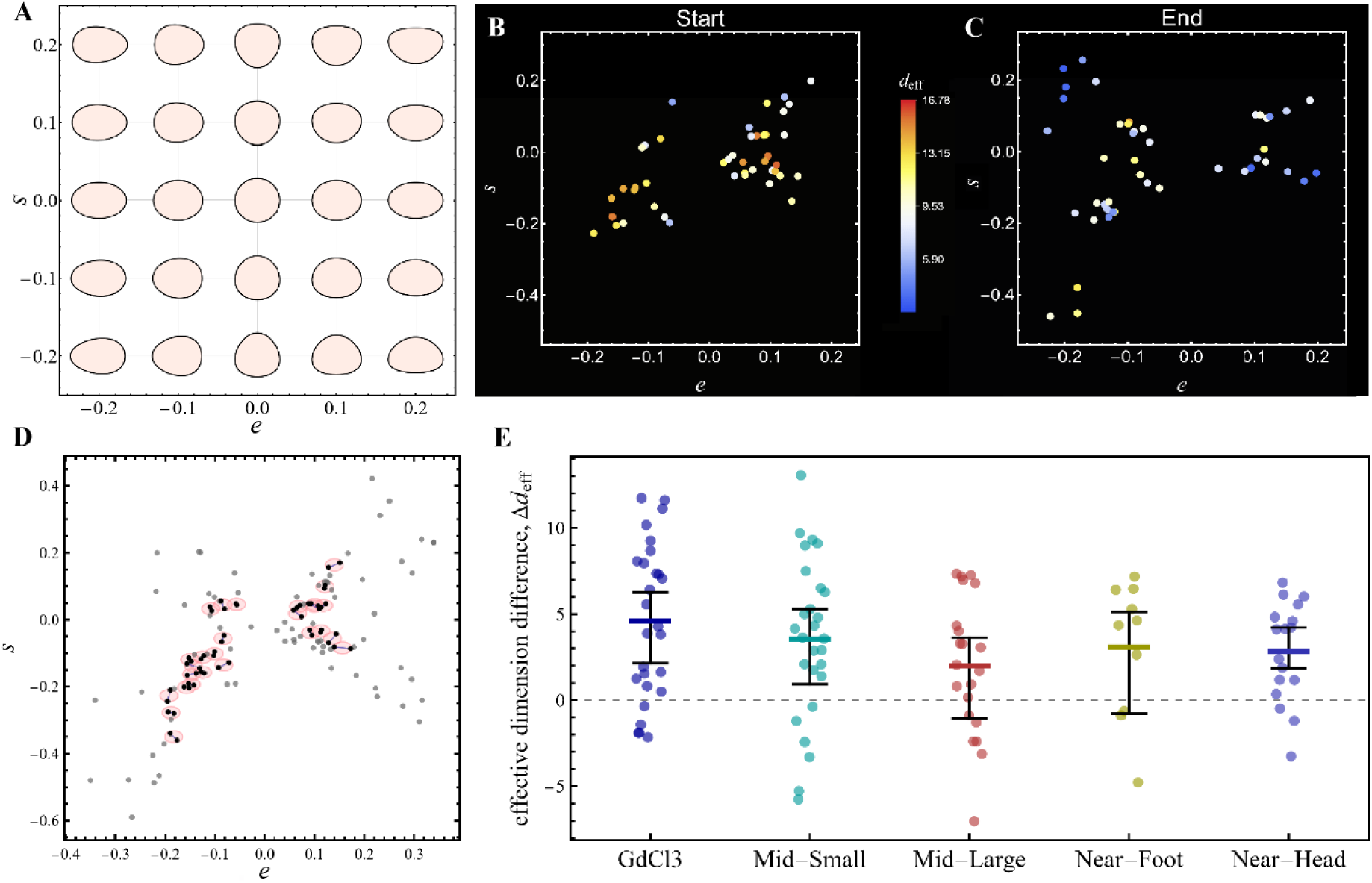
Similar coarse morphologies can correspond to different dynamical states. **(A)** Low-dimensional phenomenological shape space used to compare coarse contour morphology. The space is spanned by the parameters *e* and *s*, which describe, respectively, the leading anisotropy of the contour and a symmetric shoulder-like deviation from a purely elliptical shape (see Methods). Representative reference contours are shown in the (*e*,*s*) plane. **(B,C)** Representative maps of effective dimension, *d*_eff_, in shape space at the beginning (B) and end (C) of the preparatory period for the full dataset of GdCl₃-treated tissues. Each point represents a mean contour calculated over approximately 100 consecutive 1-min frames and plotted at its fitted location in the (*e*,*s*) plane. Color indicates the effective dimension associated with fluctuations around that local mean contour. **(D)** Shape-matching analysis for the same GdCl₃ dataset. Gray points show all available contour sets. Black points and blue links identify one-to-one matched beginning–end pairs; pink circles of radius 0.018, centered at the pair midpoints, show the matching tolerance. **(E)** Summary of the matched-shape analysis for all conditions. For each matched pair, Δ*d* = *d*^start^_eff_ − *d*^end^_eff_, so that positive values indicate a reduction in effective dimension despite approximate matching in coarse shape. Colored dots show individual pair differences descriptively and were not treated as independent biological replicates. Colored horizontal markers indicate condition means, and black error bars show 95% confidence intervals obtained by resampling tissue datasets with replacement within each condition and repeating the matching procedure.

For every matched beginning-end pair, we calculated Δ*d* = *d*^start^_eff_ − *d*^end^_eff_ (Fig. 4E). All five condition means were positive, and the condition-balanced mean was 3.21 (95% complete-tissue bootstrap CI, 1.87 to 3.86). The condition-specific confidence intervals excluded zero for GdCl₃, Mid-Small and Near-Head, but not for Mid-Large or Near-Foot, and the overall result remained stable across matching tolerances (Table S2). Because the same tissue could contribute to more than one matched pair, we also analyzed 25 pairs selected without reference to *d*_eff_, with each tissue included in at most one pair. Nineteen showed a positive difference, with a condition-balanced mean of 2.73 (95% CI, 1.43 to 3.96; exact two-sided sign-flip *P*<0.001; Fig. S9 and Table S2). Thus, the decline in effective dimension persists when coarse shape is approximately matched, supporting dynamical reorganization of the fluctuation spectrum within broadly similar morphologies rather than an effect driven solely by occasional intervals of pronounced shape change.

### Spatial calcium heterogeneity is associated with the structure of shape fluctuations

The analysis above indicates that the preparatory stage involves a hidden reorganization of morphological fluctuations. We next asked whether this organization has an identifiable physiological correlate. Calcium activity is particularly relevant because Ca²⁺-dependent excitations are coupled to muscle contraction and tissue deformation in *Hydra*^10,11,14,16,29^. We focused on the nine GdCl₃-treated tissues, for which simultaneous calcium and contour measurements were available. These tissues also maintained relatively high effective dimensions during much of the preparatory period. GdCl₃ blocks calcium channels, particularly stretch-activated channels^36,37^, including in *Hydra*^38–40^, and strongly modifies the Ca²⁺ dynamics associated with tissue rupture-repair cycles^29^.

A representative Ca²⁺ trace is shown in Fig. 5A (the remaining traces are shown in Fig. S10). Rupture events produce large Ca²⁺ transient excitations followed by slow relaxation, generating stepwise changes in the Ca²⁺ mean activity^29^. Faster spike-like activity is superimposed on this slowly varying component (see Supplementary Note 3 and Fig. S11). We therefore distinguished the interval-mean Ca²⁺ level from the faster spike intensity remaining after subtraction of the rupture-associated background (see Methods). Across rupture-defined intervals, these two quantities remained positively correlated after removal of their smooth temporal trends (Fig. 5B; equal-tissue mean *r* = 0.434, 95% CI, 0.216 to 0.657; exact two-sided sign-flip *P* = 0.0039). The result was similar after excluding the tissue represented by only four rupture-repair intervals. Thus, elevated Ca²⁺ backgrounds tend to be accompanied by stronger fast activity, although the two quantities characterize distinct temporal components of the signal.

**Fig. 5.**
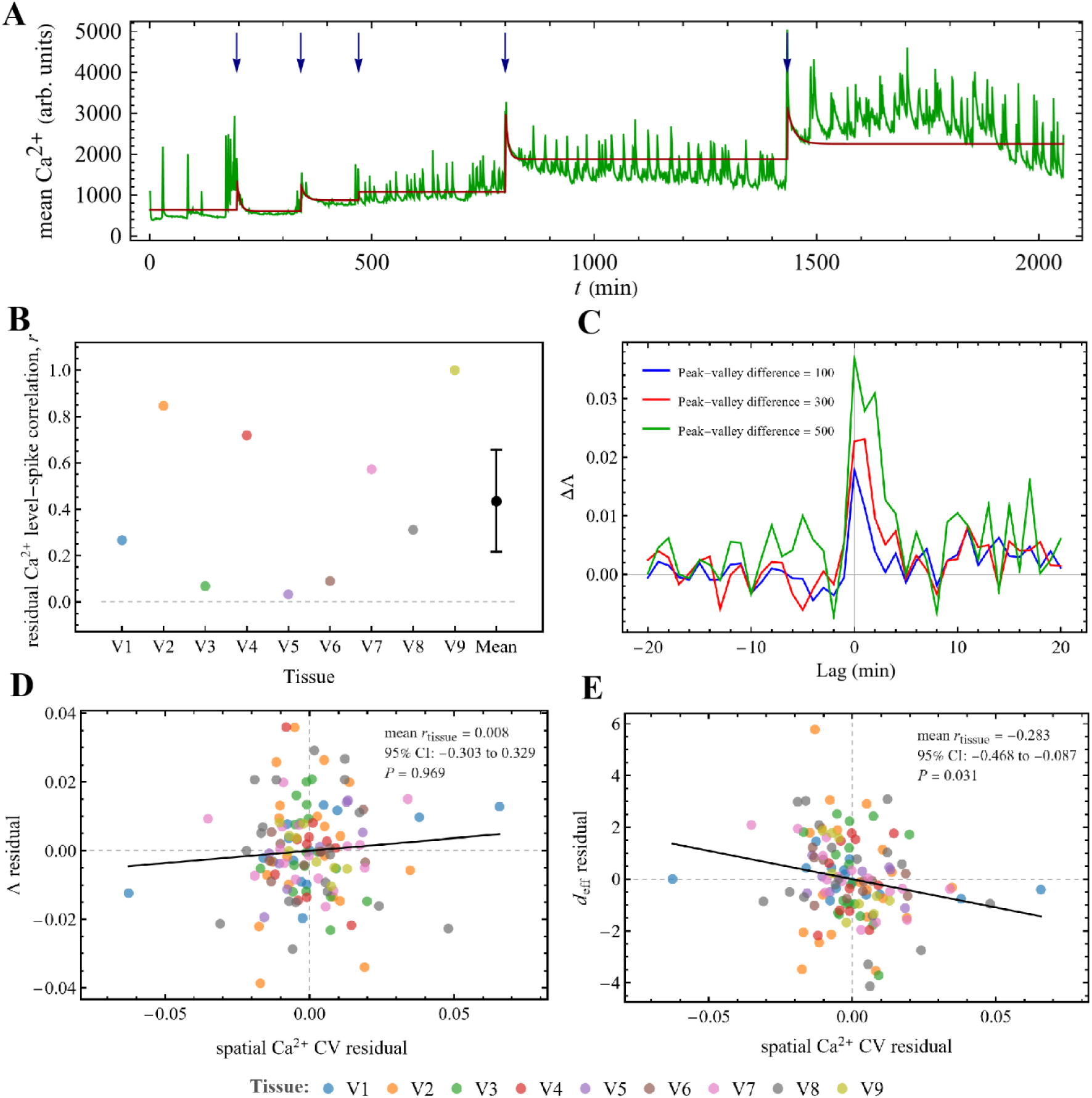
Calcium activity is associated with local shape changes and the effective dimension of shape fluctuations. (**A**) Representative mean endodermal Ca²⁺ fluorescence trace from a GdCl₃-treated tissue sample. The green curve shows the measured signal, blue arrows mark rupture events^29^, and the red curve shows the inferred rupture-associated calcium component (see Methods). (**B**) Within-tissue correlation between residual interval-mean Ca²⁺ level and residual spike intensity during the preparatory period (see Methods). Colored points represent individual tissue samples; the black point and error bar show the equal-tissue mean and its 95% bootstrap confidence interval (mean *r* = 0.434, 95% CI, 0.216 to 0.657; exact two-sided sign-flip *P* = 0.0039). (**C**) Difference, ΔΛ(τ), between event-triggered Λ profiles aligned to local maxima and minima of the Ca²⁺ signal. Tissue-specific profiles were calculated first and then averaged with equal weight across tissues. Blue, red, and green curves correspond to peak-to-valley thresholds of 100, 300, and 500 signal units, respectively. Events whose ±20-min analysis windows overlapped a rupture were excluded. All three curves show a positive peak near zero lag (exact two-sided tissue-level sign-test *P* = 0.0039, 0.039, and 0.0039, respectively). (**D**,**E**) Within-tissue associations between spatial Ca²⁺ coefficient of variation (CV) and Λ (D) or *d*_eff_ (E), after removal of temporal trends and the relevant covariates, as described in Methods. Colored points represent individual analysis windows, with tissue identity indicated by color; black lines are descriptive equal-tissue-weighted fits. No association was detected between spatial Ca²⁺ CV and Λ (mean tissue *r* = 0.008, 95% CI, −0.303 to 0.329; exact two-sided sign-flip *P* = 0.969), whereas spatial Ca²⁺ CV was negatively associated with *d*_eff_ (mean tissue *r* = −0.283, 95% CI, −0.468 to −0.087; exact two-sided sign-flip *P* = 0.031). Confidence intervals and sign-flip tests in B, D, and E use tissues as the inferential units.

We next examined whether the faster Ca²⁺ excitations component was associated with short-timescale changes in morphology. We compared event-triggered changes in the shape parameter Λ around local maxima and minima of the Ca²⁺ signal. Extrema whose ±20-min analysis windows overlapped a rupture were excluded, thereby separating these events from the large rupture-associated transients. The difference between the maximum-triggered and minimum-triggered ΔΛ profiles displayed a sharp positive peak near zero lag (Fig. 5C). This response was observed at all three peak-to-valley thresholds and increased for more prominent Ca²⁺ extrema (exact two-sided tissue-level sign-test *P* ≤ 0.039). Fast Ca²⁺ events were associated with short-timescale changes in tissue shape, consistent with the established coupling between Ca²⁺ excitation and contractile deformation in *Hydra* ^10,11,14,16,29^.

Neither the interval-mean Ca²⁺ level nor the temporal spike intensity, however, describes how the Ca²⁺ activity is distributed across the tissue. This distinction is important mechanically: a spatially uniform elevation and a spatially heterogeneous Ca²⁺ pattern can have the same mean level but produce different patterns of contractile stress and tissue deformation^10,11,14,16,29^. We therefore quantified the spatial Ca²⁺ heterogeneity using the spatial coefficient of variation (CV), a dimensionless measure of the dispersion of calcium activity across the tissue. This measure is particularly useful here because the mean Ca²⁺ signal varies strongly across tissues and following rupture events ^29^.

We first asked whether spatial Ca²⁺ heterogeneity simply reflects coarse tissue morphology. To test this, we adjusted spatial CV and Λ separately for interval-mean calcium and for smooth changes over the preparatory period, modeled as a quadratic function of normalized time (see Methods). No association was detected between the resulting residuals (Fig. 5D; equal-tissue mean *r* = 0.008, 95% CI, −0.303 to 0.329; exact two-sided sign-flip *P* = 0.969). Spatial heterogeneity in the measured calcium activity therefore did not systematically track the instantaneous coarse shape of the tissue.

By contrast, residual spatial Ca²⁺ CV was negatively associated with residual *d*_eff_ (Fig. 5E; equal-tissue mean *r* = −0.283, 95% CI, −0.468 to −0.087; exact two-sided sign-flip *P* = 0.031). Windows with more spatially heterogeneous Ca²⁺ activity thus tended to exhibit lower-dimensional, more concentrated shape fluctuations, even after accounting for time, mean Ca²⁺ level, and coarse morphology. Taken together, these analyses distinguish three aspects of Ca²⁺ activity: its overall level, its fast temporal spiking events and its spatial heterogeneity. The overall calcium level covaries with the intensity of fast activity; fast spiking events are associated with short-timescale shape changes; and spatial heterogeneity is associated with the effective dimension of shape fluctuations rather than with coarse tissue shape itself. The latter association identifies spatial calcium heterogeneity as a physiological correlate of the hidden dynamical organization revealed by *d*_eff_.

## DISCUSSION

The main result of the present study is that the prolonged quasi-spherical stage preceding the major morphological transition in regenerating *Hydra* tissue samples is not a passive quiescent period but an early phase of active dynamical reorganization manifested in the spectra of the tissue’s shape fluctuations. During this preparatory stage, the effective dimension of the shape fluctuations, *d*_eff_, decreases before any major elongation becomes apparent (Fig. 2). It shows that shape fluctuations become progressively concentrated into fewer PCA modes. This decrease is not captured by coarse morphology alone: neither the global shape parameter Λ (Fig. 3) nor comparisons at matched visible shape (Fig. 4) account for the full temporal reorganization of the dynamics. Thus, the preparatory stage is characterized by a progressive concentration of fluctuation variance into fewer dominant modes preceding the subsequent morphological transition. Since all tissue fragments examined in this study ultimately regenerated into mature normal *Hydra*^16,29^, the sample differences revealed by our analysis reflect variability in their morphogenesis trajectories rather than in the final morphogenesis outcome.

Positional cues inherited from the parent *Hydra* are reflected in the fluctuation dynamics prior to significant morphological changes. Tissue samples retaining positional information (i.e., large central tissue fragments or those excised near the head or the foot) tend to reach lower effective dimensions during the preparatory period (Fig. 2). This conclusion is consistent with our previous finding that excised *Hydra* fragments retain positional memory from the parent animal and that this memory is reflected in their morphological dynamics ^16^. In addition, a spatial gradient in the Ca²⁺ activity is aligned with the future polarity axis from the onset of morphogenesis, before major morphological changes become apparent^16^. This gradient may contribute to concentrating shape fluctuations into fewer dominant directions. GdCl₃-treated tissue samples, in which the ability of Ca²⁺ activity to respond to morphological changes is weakened^29^, maintain higher effective dimension and a less concentrated fluctuation spectrum.

These findings suggest a broader physical interpretation of canalized morphogenesis. In our earlier work, morphogenesis was described in terms of an effective morphological potential that evolves slowly in time, such that its exponential defines the probability density for finding the tissue near a given morphological configuration^13,14^. In the simplest picture, this potential has a double-well structure, with one minimum corresponding to the approximately spherical tissue state and the other to the elongated state characteristic of the *Hydra*’s mature body axis. The main morphological transition then occurs through an activation-like process, once the elongated minimum becomes sufficiently favorable. This is possible due to global tilting dynamics of the morphological potential. Positional cues lead to a more complex effective potential, consistent with more continuous transitions and, in some cases, with passage through quasi-metastable intermediate states^16^. In our phenomenological description, shape fluctuations, for which stochastic Ca²⁺ activity is a principal source of active noise, evolve much faster than the effective morphological potential. Over each PCA interval analyzed here, the potential is therefore approximately stationary, and the fluctuations sample a quasi-stationary distribution around its quasi-spherical minimum. The covariance spectrum probes the local structure of this effective landscape. Because the fluctuations are actively driven, however, the PCA eigenvalues reflect the landscape curvature together with the statistics of the active drive, rather than mechanical stiffness alone. Across successive intervals, the leading eigenvalue grows disproportionately and its direction shows modest temporal stabilization (Fig. S7). Thus, the total fluctuation variance increases, and its distribution becomes increasingly anisotropic and concentrated along the leading PCA eigenvector direction. This increasing dominance of the leading mode is the principal spectral driver of the decline in *d*_eff_. Within the effective-potential framework, these results support a picture of progressive reshaping of the morphological potential before significant shape change. This progressive concentration and temporal organization of morphological fluctuations is what we identify operationally as dynamical canalization.

The calcium analysis of GdCl₃-treated tissues provides a mechanophysical viewpoint of the observed spectral concentration. A regenerating *Hydra* fragment is a pressurized epitheliomuscular shell whose shape is controlled by calcium-dependent actomyosin contraction acting together with hydrostatic pressure^14,16^. What matters for shape change is not only the overall level of Ca^2+^ activity, but also how unevenly that activity is distributed across the tissue. Spatial differences in calcium-dependent contraction can generate uneven stresses that, together with lumen pressure, preferentially excite particular modes of deformation. Consistent with this picture, spatial Ca^2+^ coefficient of variation was negatively associated with *d*_eff_ after accounting for within-tissue temporal trends, interval-mean calcium and coarse morphology. No association with Λ was detected after accounting for temporal trends and interval-mean calcium. Thus, intervals with greater spatial heterogeneity of Ca^2+^ activity tended to exhibit stronger concentration of fluctuation variance into fewer dominant modes. This association is consistent with spatial differences in calcium-dependent contractile activity contributing to the emergence of dominant deformation modes during the preparatory stage.

The comparison with untreated tissue samples addresses a different aspect of this relation. GdCl₃-treated samples occupied a higher-dimensional fluctuation regime during much of the preparatory period, although *d*_eff_ also declined progressively for these samples. Because GdCl₃ interferes with mechanosensitive calcium signaling ^29,37–40^, the shift toward higher *d*_eff_ is consistent with such signaling influencing the organization of the fluctuation spectrum. The treated tissue samples therefore exhibit both a systematic shift toward higher dimensionality and a continued temporal concentration of the fluctuation spectrum.

The calcium-dependent fluctuation spectra suggest a possible mechanophysical link between spatially heterogeneous Ca^2+^ activity and the observed spectral concentration. In regenerating *Hydra*, this spectral concentration is the defining signature of dynamical canalization: as total fluctuation variance grows, the fluctuation spectrum becomes increasingly dominated by a smaller set of deformation modes. More generally, our findings indicate that morphogenetic organization can become evident in fluctuation dynamics before it appears in coarse tissue morphology. Following the evolution of fluctuation spectra may therefore reveal similar preparatory processes in other developing systems.

## MATERIALS AND METHODS

### Experimental Methods

All experiments were carried out with a transgenic strain of *Hydra vulgaris* (*AEP*) carrying a GCaMP6s fluorescence probe for Ca^2+^ (see Refs.^10,11,14^ for details of the strain). Animals were cultivated in a *Hydra* culture medium (HM; 1mM NaHCO3, 1mM CaCl2, 0.1mM MgCl2, 0.1mM KCl, 1mM Tris-HCl pH 7.7) at 18°C. The animals were fed every other day with live *Artemia nauplii* and washed after ∼4 hours. Experiments were initiated ∼24 hours after feeding. Small tissue fragments were excised from different regions along the axis of a mature *Hydra*. These tissue fragments were incubated in a dish for ∼3 hrs to allow their folding into spheroids prior to transferring them to the experimental sample holder under flow of HM.

For the experiments with GdCl_3_: 1M stock (Sigma 439770 powder dissolved in water) was diluted to a final concentration of 50μM in HM medium and introduced to the experimental setup under the same flow conditions throughout the experiment as above.

The experimental setup was similar to the one described before^16^. In all the experiments, spheroid tissues were placed within wells of ∼1.3 mm diameter made in a strip of 2% agarose gel (Sigma) to keep the regenerating *Hydra* in place during time-lapse imaging. The tissue spheroids were free to move within the wells. All the experiments were done at room temperature with a continuous medium flow as described before^16^.

### Microscopy

Time-lapse bright-field and fluorescence images were taken by a Zeiss Axio-observer microscope (Zeiss, Oberkochen Germany) with a 5× air objective (NA=0.25) and a 1.6× optovar and acquired by a CCD camera (Zyla 5.5 sCMOS, Andor, Belfast, Northern Ireland). The sample holder was placed on a movable stage (Marzhauser, Germany), and the entire microscopy system was operated by Micromanager, recording images at 1 min intervals. The recording at 1 min resolution allowed faithful analysis of the area drops over long experiments, without tissue damage, while also enabling recording from multiple tissue samples.

### Data Analysis

For the analysis, images were reduced to 696−520 pixels (∼1.6 µm per pixel) using ImageJ. Masks depicting the projected tissue shape were determined for a time-lapse movie using the bright-field (BF) images by a segmentation algorithm described in ^41^ and a custom code written in Matlab. Shape analysis of regenerating *Hydra*’s tissue was done by representing the projected shape of the tissue by polygonal outlines as presented before^42^. The fluorescence analysis was done on images reduced to the same size as the bright-field ones (696−520 pixels).

### Contour preprocessing and alignment before PCA

Projected tissue contours were extracted frame by frame and represented as ordered sets of boundary 100 polygon vertices in the image plane. Before principal component analysis, all contours were transformed into a common shape-centered coordinate system so that the PCA would capture changes in contour geometry rather than trivial differences in size, position, orientation, or boundary parametrization. The centroid and area were computed using the standard polygon formulas applied to the contour points.

Each contour was first rescaled so that the enclosed projected area was equal to unity. This normalization removed global size changes, in particular those associated with inflation of the tissue lumen and allowed the subsequent analysis to focus specifically on shape fluctuations. The contour was then translated so that its centroid lay at the origin.

To ensure consistent orientation, contours were represented with a fixed traversal direction by enforcing counterclockwise ordering of vertex points according to the sign of the polygon signed area. Candidate rotation angles were then estimated from the contour geometry using both the principal axis of the covariance matrix and the diameter defined by the most distant pair of vertex points. Because these estimates are subject to π/2 and π ambiguities, multiple candidate orientations were evaluated. For each frame, the selected orientation minimized a cost function combining three terms: preference for horizontal alignment of the main axis, agreement with the recent contour history, and a temporal smoothness penalty suppressing abrupt 180° flips between consecutive frames.

After rotational alignment, residual differences in contour phase were removed by cyclic reindexing of the contour points. When previous aligned contours were available, the optimal cyclic shift was chosen so as to minimize mismatch with a weighted history of recent contours. For the first contour, an anchor point was selected deterministically. The result of this procedure was a time series of contours expressed in a common reference frame, with unit area, centroid at the origin, consistent orientation, and consistent contour phase. PCA was then applied to these aligned contours within each selected time window. In this way, the resulting principal modes reflect genuine changes in contour shape and fluctuation structure rather than trivial variation due to inflation, translation, rotation, or contour indexing.

### PCA of shape fluctuations

Projected contours were represented as ordered arrays of Cartesian coordinates, preprocessed as described above. To obtain a fixed-length representation suitable for PCA, each normalized contour was converted into a radial signature *r* (*θ*). Specifically, the polar angle was divided into *m* equal bins (here *m* =100), and for each angular bin the maximal radial distance of the contour points falling in that sector was recorded. The bins were always ordered by increasing angle from a fixed reference direction, spanning the interval [−π,π), so that the same component of the signature always corresponded to the same angular sector in all contours. Bins containing no contour point were filled by linear interpolation on the circular domain. The resulting radial profile was then normalized by its mean radius, yielding a scale-independent one-dimensional descriptor of shape. Thus, each contour was represented by a vector of length 100.

PCA was then performed directly on the matrix of radial signatures. Denoting by *r_i_(j)* the *i*-th radial-signature vector of contour *j*, we first computed the mean signature

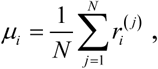

where *N* is the number of contours included within one time window. We then constructed the centered data matrix

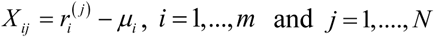

and the covariance matrix of the centered signatures, *C* = *XX ^T^* (*N* −1). PCA mode *k* is then defined as the eigenvector ***v***^(*k*)^ satisfying *C**v***^(*k*)^ = *λ* ***v***^(*k*)^, with modes ordered by decreasing *λ*_k_. The projection coefficient of contour *j* onto mode *k* was *a^(k)^_j_ = Σ_i_v^(k)^_i_X_ij_*. For visualization of the principal modes, the mean radial signature was perturbed along each eigenvector according to ***μ*** ± *b**v***^(*k*)^, where *b* is a display scale chosen for best visualization. These perturbed signatures were then mapped back into Cartesian coordinates, thereby producing the deformed contours shown for the positive and negative directions of each PCA mode. This representation allows one to interpret each principal component directly as a mode of contour deformation about the mean shape.

To assess sensitivity to analysis choices, the PCA and effective-dimension calculation were repeated using radial signatures containing 50, 75 or 100 angular bins, using either harmonic-based or centroid-based centering, and after temporal subsampling at 5, 10 or 20 min intervals. Initial-to-final changes were calculated for each tissue and summarized with equal weighting of the five experimental conditions (Fig. S6).

### Shape Space for coarse contour matching

To compare tissues with similar coarse morphologies, we introduced a low-dimensional phenomenological shape space based on a fitted family of normalized closed contours. After the preprocessing described above, each contour was represented in complex form, resampled uniformly along arc length, centered, and normalized to unit enclosed area. A radial signature was then computed by dividing the angle into *m* bins and assigning to each bin the maximal radius of the contour in that angular sector, followed by normalization by the mean radius. This yielded a scale-independent one-dimensional representation of contour shape. The reference family was defined by a two-parameter set of smooth closed contours, described by continuous parameters *e* and *s*, using the parametric representation:

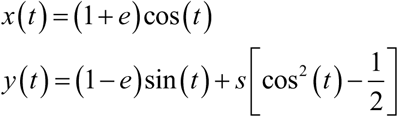

The parameter *e* captures the leading anisotropy of the contour, whereas *s* captures a symmetric shoulder-like deviation from a purely elliptical shape. In addition, two related branches of this family were considered, denoted *idx* = 0 and *idx* = 1, where the second branch is obtained from the first by a 90^0^ rotation.

For each target contour, the fit was performed separately for the two branches. In each case, the squared distance between the target radial signature and the radial signature of the reference family was minimized numerically, using a coarse grid in (*e,s*) for initialization followed by local refinement. The final assignment of the contour in shape space was taken to be the better fitted of the two, namely the one with the smaller residual distance. Thus, each contour was assigned effective shape coordinates (*e,s*) together with the branch index of the best-fitting family.

Because the contours had already been pre-aligned so that their principal axis was typically horizontal, the shapes shown in Fig. 4A were displayed using the *idx* = 0 branch for *e* ≥ 0 and the *idx* = 1 branch for *e* < 0, which correspond to the dominant orientations observed in the data. The resulting (*e,s*) space provides a compact representation of coarse contour geometry: contours mapped to nearby points are similar in overall form, even if they differ in finer details. This makes the space useful for identifying approximately shape-matched tissues and for testing whether effective dimension changes systematically even when coarse morphology is held approximately fixed.

### Extrema detection from calcium fluorescence traces

To identify the principal extrema of the Ca^2+^ fluorescence signal, fluorescence traces were first smoothed with a Gaussian kernel of width σ = 2 frames. Local maxima and minima were then identified on the smoothed trace from changes in the sign of the first derivative. Prominent extrema were selected by requiring a minimum peak-to-valley difference of 100 signal units. This threshold was chosen to capture the most significant extrema while avoiding an excessive number of detected points. The event-triggered analysis was repeated using peak-to-valley thresholds of 100, 300 and 500 signal units.

### Construction of the rupture-associated calcium component

To estimate the rupture-associated calcium component, each calcium fluorescence trace was divided into successive intervals defined by the rupture-time list (see, e.g., the blue vertical lines in Fig. 5A). Within each interval, the signal was fitted by a constant plus a decaying exponential. The fit was constrained by the prominent minima identified as described above: the exponential baseline was chosen so as to pass close to the minima points within that interval, thereby using them as anchor points for the relaxing background. This procedure was intended to capture the slow relaxation that follows each rupture event while excluding the superimposed spike-like excursions. The fits from successive intervals were then combined to produce a full trace, interpreted operationally as the rupture-associated calcium component. The red curve in Fig. 5A shows one representative example.

### Event-triggered calcium-shape analysis

The event-triggered analysis was performed using peak-to-valley thresholds of 100, 300 and 500 signal units. Extrema whose ±20-min analysis windows overlapped a rupture event were excluded. Within each tissue, event-triggered Λ profiles were averaged separately around the calcium maxima and minima. Their difference was defined as ΔΛ(τ) = ⟨Λ(*t*_max_ + τ)⟩ − ⟨Λ(*t*_min_ + τ)⟩, where τ is the lag relative to the calcium extremum. Tissue-specific profiles were calculated first and then averaged with equal weight across tissues. The zero-lag effects were evaluated using exact two-sided tissue-level sign tests.

### Within-tissue residual correlations

The associations shown in Fig. 5B,D,E were calculated separately for each tissue. For every comparison, the corresponding two variables were first regressed separately on the relevant control variables, and the Pearson correlation was then calculated between their residuals. In Fig. 5B, interval-mean calcium and spike intensity were each adjusted for a quadratic temporal trend in normalized time. For Fig. 5D, spatial calcium CV and Λ were each regressed on interval-mean calcium and normalized time, with both linear and quadratic time terms included. For Fig. 5E, spatial calcium CV and *d*_eff_ were each regressed on interval-mean calcium, Λ and normalized time, again with both linear and quadratic time terms included. Thus, the analysis in Fig. 5E measures the association between spatial calcium heterogeneity and effective dimension that is not accounted for by their common temporal evolution, mean calcium level or coarse tissue shape.

Each comparison yielded one correlation coefficient per tissue. The nine tissue-level coefficients were averaged with equal weight, preventing tissues with more intervals or analysis windows from contributing disproportionately. Confidence intervals for the mean correlation were obtained by bootstrap resampling of tissues. Exact two-sided P values were calculated by comparing the observed mean correlation with the distribution obtained from all 2⁹ possible sign reversals of the nine tissue-level coefficients. For Fig. 5B, the analysis was also repeated after excluding the tissue containing only four rupture-defined intervals.

### Use of artificial intelligence tools

ChatGPT (OpenAI) and Claude (Anthropic) were used to assist with language editing and with the development and verification of analysis code. All code, numerical results and scientific interpretations were reviewed and verified by the authors.

## Author Contributions

E.B. and O.A. designed the experiments and analyzed the data. E.B. conducted the experiments. O.A. statistically analyzed the data. E.B. and O.A. wrote the manuscript.

## Data and resource availability

All relevant data and resource details are provided in the article and its Supplementary Information. Analysis code and processed contour data are available from the corresponding authors upon reasonable request.

## Supplementary Information for

**Table S1.** Dataset provenance and acquisition characteristics.

| Condition | n | Dataset provenance and source | Preparatory-stage duration, h<br>median (range) | Ca<sup>2+</sup><br>fluorescence |
| --- | --- | --- | --- | --- |
| Mid-Small | 9 | Six tissues from the phase-transition study (Ref. 14);<br>three additional tissues acquired during revision of the<br>related Biophysical Journal study | 27.1 (16.0-40.0) | Recorded; not<br>analyzed here |
| Mid-Large | 14 | Positional-memory study (Ref. 16) | 14.7 (10.0-31.6) | Recorded; not<br>analyzed here |
| Near-Head | 8 | Positional-memory study (Ref. 16) | 26.6 (17.5-31.5) | Recorded; not<br>analyzed here |
| Near-Foot | 8 | Positional-memory study (Ref. 16) | 25.0 (11.0-35.0) | Recorded; not<br>analyzed here |
| GdCl<sub>3</sub> | 9 | New recordings; also analyzed in the parallel rupture-repair study (Ref. 29) | 24.7 (16.6-39.0) | Recorded and<br>analyzed here |

**Note:** The preparatory-stage duration was defined as the elapsed time between the first and last frames included in the preparatory-stage analysis. Values (in hrs) are the median, with the full range in parentheses. All tissues were recorded at 1-min intervals using simultaneous bright-field and GCaMP6s fluorescence imaging. Calcium-specific analyses in the present manuscript were restricted to the GdCl<sub>3</sub>-treated tissues.

**Fig. S1.**
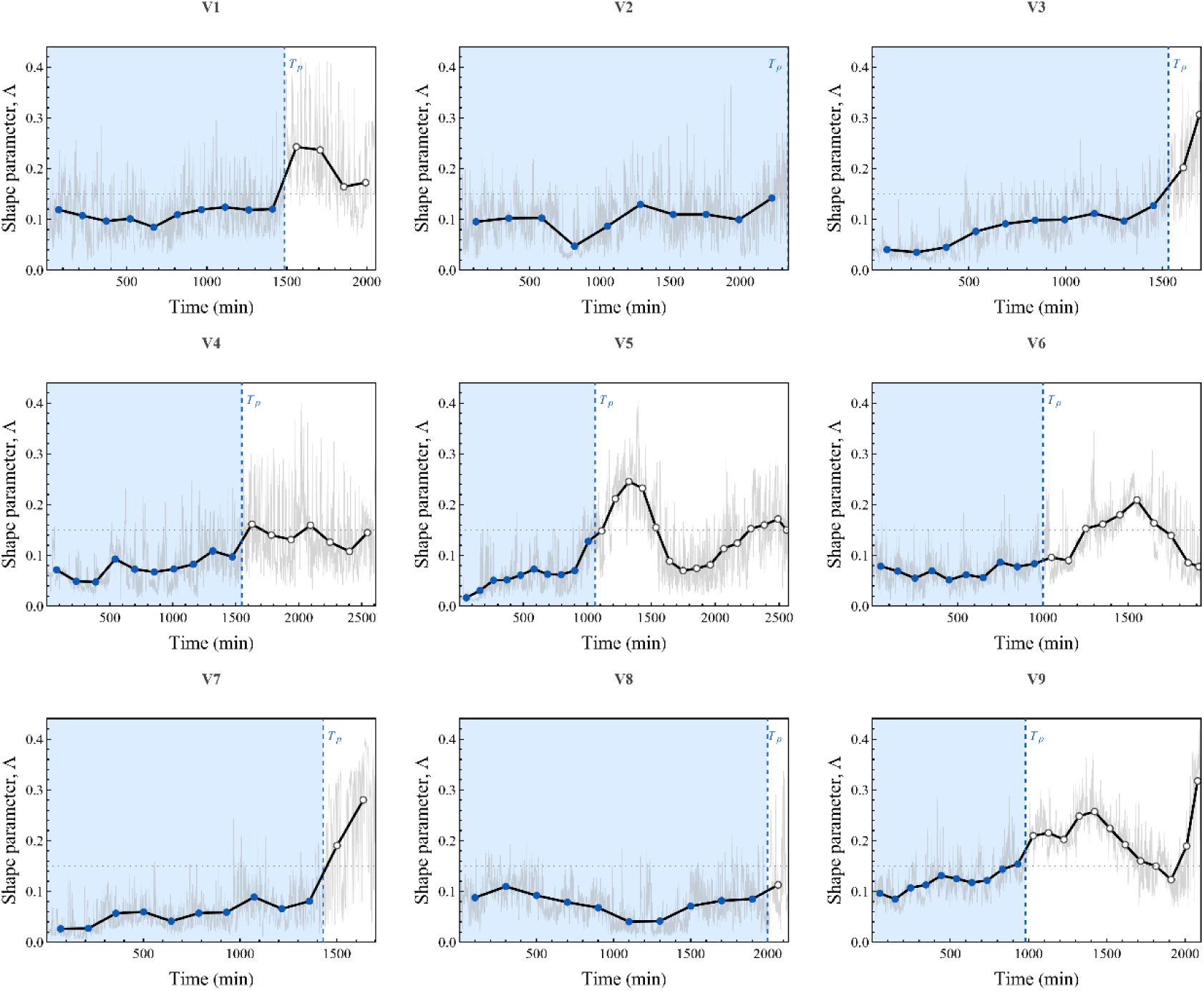
Complete shape-parameter trajectories of the GdCl₃-treated tissues. Pale-gray curves show the framewise shape parameter Λ for each tissue. Black curves and symbols show averages over consecutive non-overlapping windows of duration *T*_p_/10, where *T*_p_ is the tissue-specific duration of the early preparatory period. Filled blue symbols and blue shading indicate the analyzed period; open symbols show later intervals. Dashed vertical lines mark *T*_p_, and the dotted horizontal line marks Λ=0.15.

### Supplementary Note 1: Robustness of the effective dimension to rupture-associated volume drops

During the early stages of *Hydra* regeneration, tissue fragments undergo repeated inflation–rupture– repair cycles, which appear in the projected-area trace as a characteristic sawtooth pattern: slow swelling punctuated by abrupt drops associated with pressure release[1-3]. In the main text, the effective dimension *d*_eff_ is computed from PCA of preprocessed contour shapes during the preparatory stage, after rescaling each contour to unit area and aligning it in a common reference frame. The purpose of this Supplementary Note is to examine whether the intermittent volume-drop events intrinsic to this stage can bias that calculation.

We address this question in three steps. First, we quantify the contour-shape changes associated with individual drop events and compare them with the changes accumulated during the preceding inflation segments. Second, we recompute *d*_eff_ using two alternative segmentations of the same preparatory-stage data: the equal normalized time intervals used in the main text, and an alternative partition defined by successive drops, excluding the drop frames themselves. Third, we interpret the outcome in PCA terms by asking whether drop events mainly contribute fluctuation amplitude or instead introduce additional fluctuation directions that would substantially alter the effective dimensionality of the contour dynamics.

#### Within-sawtooth and across-drop contour-shape changes

We begin with the first step. Figure S2A shows a representative projected-area trace during the preparatory stage. The trace displays the characteristic sawtooth dynamics of gradual inflation punctuated by abrupt drops. On this trace, A, B, and C mark the centers of three short contour-averaging windows used to define the geometric comparison. Each representative contour was obtained by averaging over 6 consecutive frames (6 min). Contour-shape change was quantified as the Euclidean distance between the 100-component, mean-radius-normalized radial signatures of the two area-normalized and aligned averaged contours (see Methods). The A and B windows lie within the same sawtooth interval, with the upper end of the B window located 1 frame (1 min) before the rupture point, whereas the C window lies after the drop, with its lower end 1 frame after the rupture point. The shape difference between the averaged contours near A and B was taken as the within-sawtooth change, whereas the difference between the averaged contours near B and C was taken as the across-drop change.

The results are summarized in Figs. S2B,C. Across all five tissue classes, the two types of contour-shape change are of the same order of magnitude, showing that rupture events are not geometrically negligible in the normalized contour representation. At the same time, within-sawtooth changes are typically somewhat larger than across-drop changes. The tissue-level analysis confirmed this trend: for 44 of the 48 tissues contributing valid rupture events, the median paired difference between the within-sawtooth and across-drop distances was positive. The mean tissue-level difference was also positive in all five conditions.

This comparison establishes an important point for the subsequent analysis. The drop events do introduce substantial contour-shape changes and therefore cannot simply be ignored. However, they do not dominate the contour dynamics during the preparatory stage. A substantial part of the geometric evolution is accumulated within the sawtooth intervals between drops. This motivates the next step of the analysis, in which we test directly whether the temporal behavior of *d*_eff_ depends on the way the preparatory stage is partitioned: either into equal normalized time intervals or into intervals defined relative to the drops.

**Fig. S2.**
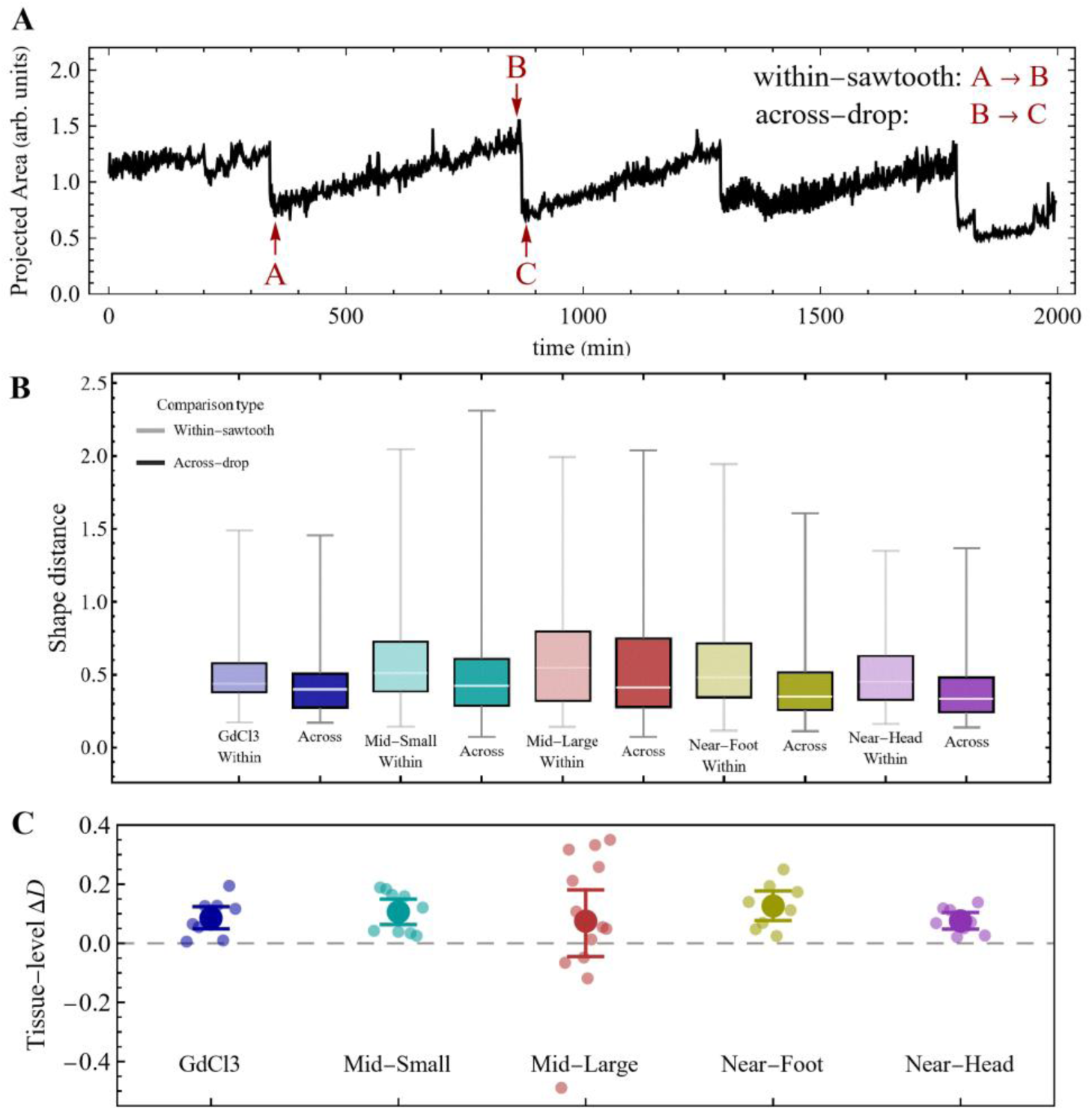
Contour-shape changes within sawtooth intervals and across volume-drop events. (A) Representative projected-area trace during the preparatory stage, showing gradual inflation interrupted by abrupt drops. A, B, and C mark the centers of the contour-averaging windows used for the comparison. Each representative contour was averaged over six consecutive frames. The A and B windows lie within the same sawtooth interval, with the B window ending one frame before the rupture point. The C window begins one frame after the rupture. The contour difference from A to B defines the within-sawtooth distance, whereas that from B to C defines the across-drop distance. (B) Pooled event-level distributions of within-sawtooth and across-drop contour changes for the five experimental conditions. These distributions are shown descriptively because multiple events originate from the same tissue. Lighter boxes denote within-sawtooth distances and darker boxes denote across-drop distances. (C) Tissue-level paired comparison. For each tissue, Δ*D* is the median across rupture events of the paired within-sawtooth distance minus across-drop distance. Small symbols show individual tissues; large colored symbols and error bars show condition means and tissue-bootstrap 95% confidence intervals. The within-sawtooth distance exceeded the across-drop distance in 44 of 48 tissues, and the mean difference was positive in each of the five conditions. The equal-condition mean was 0.0936 (condition-stratified bootstrap 95% CI, 0.0650–0.1207; two-sided tissue-level sign-flip p < 10^-5^).

#### Robustness of the effective dimension to segmentation relative to volume-drop events

We next check whether the temporal decline in *d*_eff_ depends on how the preparatory stage is partitioned relative to the volume-drop events. In the main text, each preparatory-stage trajectory was normalized in duration, divided into 10 equal intervals, and analyzed by PCA separately in each interval.

To test the influence of the drop timing, we repeated the same PCA-based calculation using an alternative segmentation defined by the drop events themselves. For each sample, the list of drop times was used to partition the preparatory stage into intervals between successive drops, with the drop frames excluded from the analysis. PCA was then performed separately on the set of contours belonging to each such inter-drop interval, and the corresponding *d*_eff_ values were assigned to the normalized time coordinate given by the center of that interval. Because different samples contain different numbers of drops and therefore different numbers of inter-drop intervals, the resulting per-sample curves do not share a common set of time points. To compare them across samples, the drop-defined curves were interpolated onto a common normalized grid and then averaged within each tissue class. By contrast, in the equal-interval analysis, all samples contribute directly to the same 10 interval indices, so the class mean is obtained simply by pointwise averaging across samples.

**Fig. S3.**
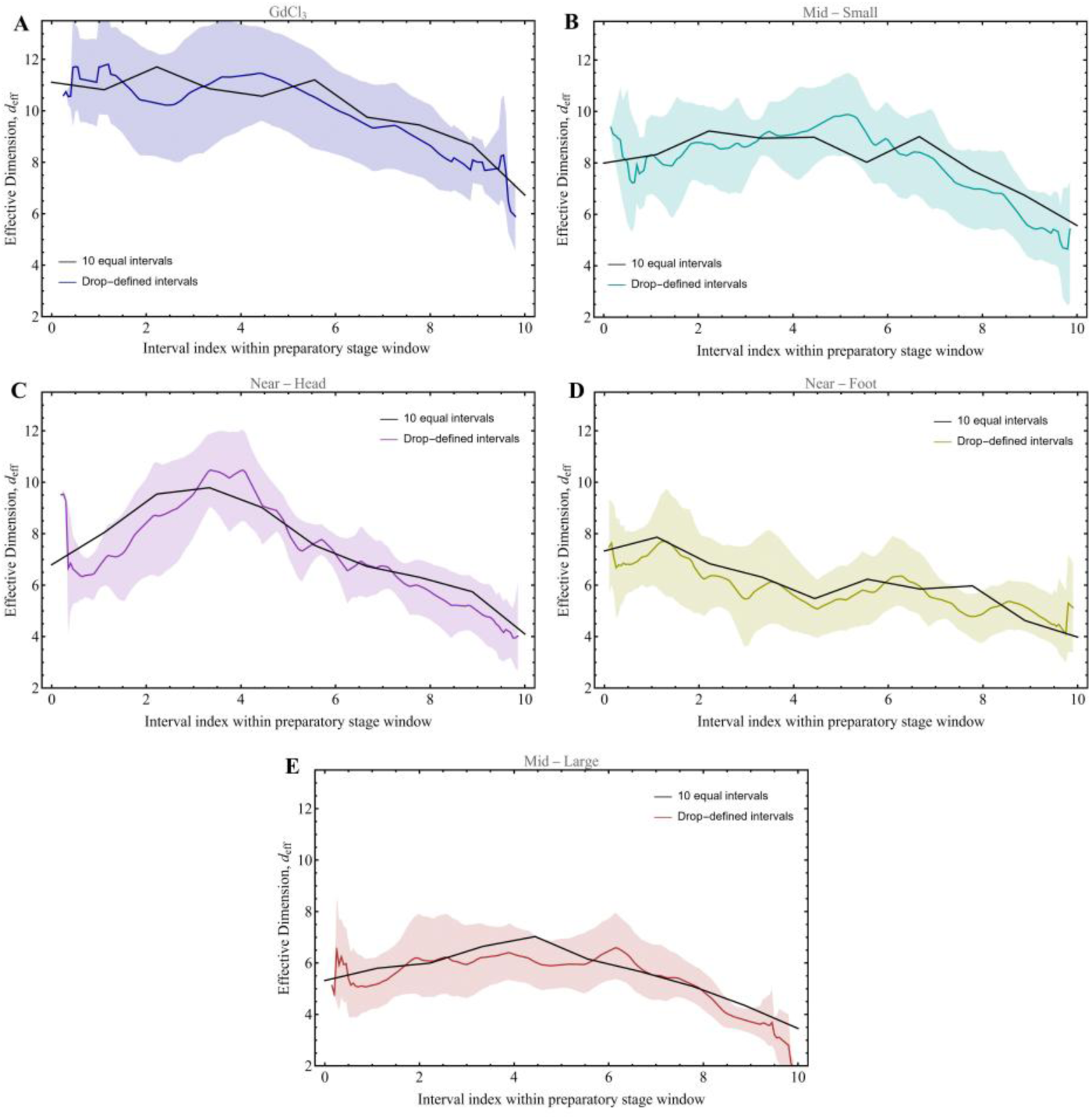
Effective dimension from equal and drop-defined intervals. (A–E) Comparison of the temporal evolution of the effective dimension deff during the preparatory stage for the five tissue classes: **(A)** GdCl_3_, **(B)** Mid-Small, **(C)** Near-Head, **(D)** Near-Foot, and **(E)** Mid-Large. In each panel, the black curve shows the class-averaged *d*_eff_ obtained by dividing the preparatory stage into 10 equal normalized time intervals, as in the main text. The colored curve shows the corresponding class-averaged *d*_eff_ obtained from intervals between successive drop events, with the drop frames excluded. For the drop-defined analysis, the colored shaded region indicates the bootstrap confidence band around the mean curve. Although the drop-defined curves are noisier, reflecting the irregular duration and number of inter-drop intervals across samples, the same overall temporal decrease in *d*_eff_ is recovered in all tissue classes. This shows that the decline in effective dimension during the preparatory stage is not an artifact of the equal-interval partition used in the main text.

The results are shown in Fig. S3. In all five tissue classes, the drop-defined calculation yields the same qualitative behavior seen in the main-text analysis: *d*_eff_ is generally higher at early times and lower toward the end of the preparatory stage. Thus, the progressive reduction in effective dimensions does not disappear when the analysis is reformulated in a way that explicitly respects the timing of the drop events. This indicates that the decrease in *d*_eff_ is not an artifact of the equal-interval partition used in the main text, but a robust feature of the underlying dynamics.

At the same time, the drop-defined curves are visibly noisier than the curves obtained from the 10 equal intervals. This increased noise is expected from the way the alternative analysis is constructed. First, the inter-drop intervals vary substantially in duration, so the PCA in a given interval may be based on very different numbers of contours from one interval to the next; short intervals naturally provide less stable estimates of the covariance structure and hence of *d*_eff_. Second, the number of inter-drop intervals differs from sample to sample, unlike the fixed set of 10 equal intervals used in the main analysis. As a result, the drop-defined class averages must be assembled from irregularly spaced per-sample estimates after interpolation to a common normalized time axis, which introduces additional variability. Third, because the drop-defined intervals are aligned to event timing rather than to a fixed global partition, neighboring points on the averaged curve need not correspond to the same number of contributing samples or to intervals of comparable temporal width. Together, these features make the drop-defined estimate intrinsically less smooth, even when the underlying biological trend is the same.

The important point, however, is that the extra noise does not alter the main conclusion. In every tissue class, the drop-defined analysis remains broadly consistent with the equal-interval analysis, showing that the reduction in effective dimension during the preparatory stage is not driven by the particular placement of fixed analysis windows relative to rupture events. Rather, it reflects a genuine temporal reorganization of the fluctuation dynamics that persists even when the analysis is reformulated in a way that tracks the intermittent sawtooth structure of the projected-area trace.

#### Why rupture events do not strongly alter the effective dimension

Figs. S2 and S3 suggest an interesting picture. On the one hand, rupture events are not negligible at the level of contour geometry: as shown in Fig. S2, they are associated with shape changes that are comparable in magnitude to those accumulated during the inflation intervals between successive drops. On the other hand, when the effective dimension is recomputed using intervals defined by the drops themselves, the resulting temporal profiles remain similar to those obtained from the 10 equal intervals used in the main text (Fig. S3). Thus, rupture events can produce substantial shape displacements without strongly altering the effective dimensionality of the fluctuations.

To understand this seemingly paradoxical result, it is important to recall that *d*_eff_ does not quantify the magnitude of a shape change itself. Rather, it reflects how shape-fluctuation variance is distributed across PCA modes. A rupture event can therefore produce a substantial displacement in shape space while still leaving the dominant fluctuation directions largely unchanged. In such a situation, the event mainly adds amplitude along modes that are already present, instead of generating a qualitatively new set of independent fluctuation directions. From the PCA point of view, a rupture event can perturb the shape trajectory without fundamentally reorganizing the dominant fluctuation directions

If this interpretation is correct, it suggests that the dominant fluctuation modes are relatively robust even to abrupt rupture–repair episodes. This, in turn, implies that the progressive reduction in effective dimension during the preparatory stage is not simply reset by individual rupture events. Rather, the tissue may be undergoing a slower internal reorganization that persists through these events and progressively concentrates shape-fluctuation variance into fewer dominant PCA directions. In this sense, the robustness of *d*_eff_ to drop-defined segmentation is consistent with the possibility that the observed decrease in effective dimension reflects hidden internal changes taking place during the preparatory stage, which continue to shape the dominant fluctuation structure even in the presence of abrupt rupture events.

**Fig. S4.**
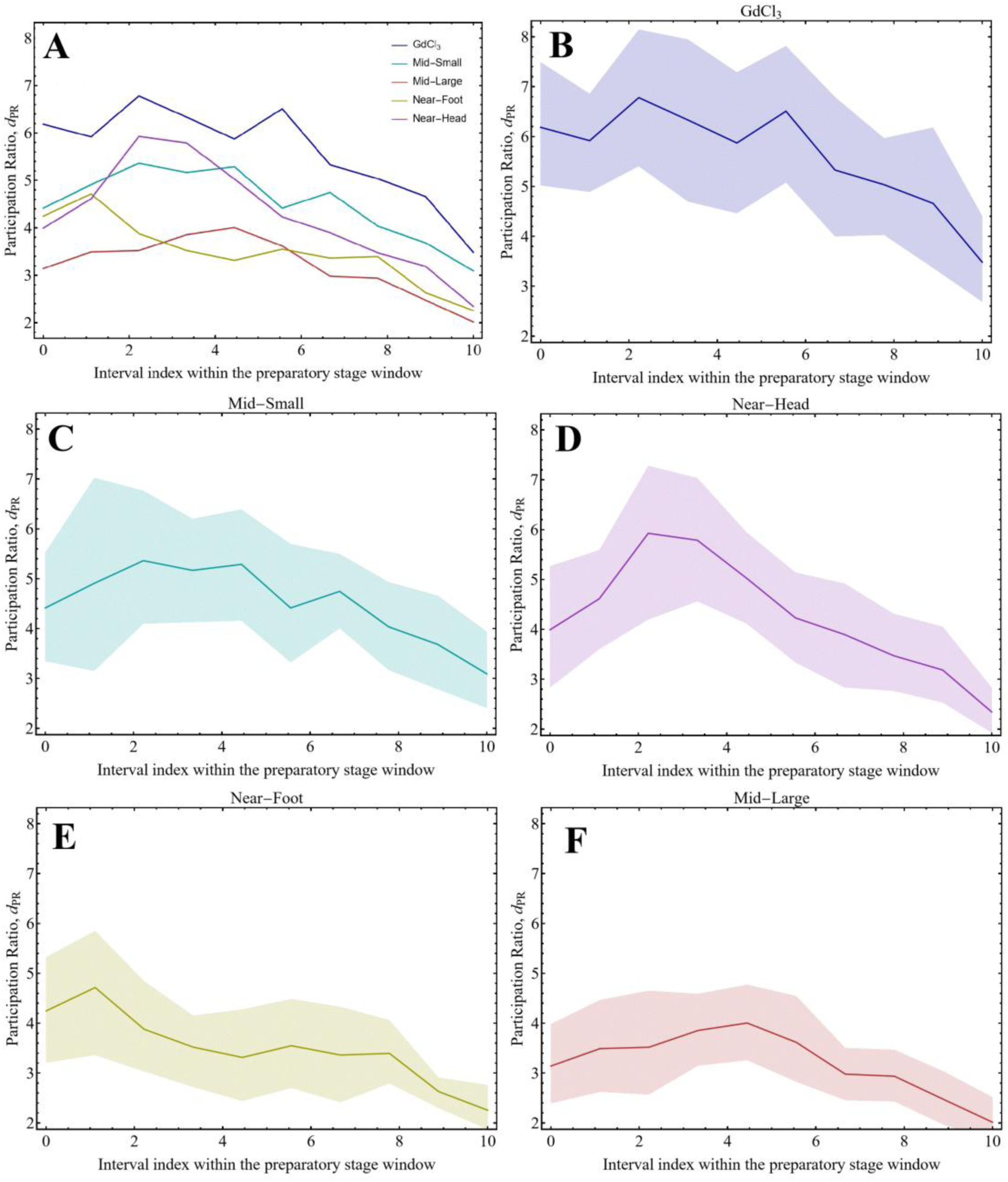
Participation-ratio measure of fluctuation dimensionality during the preparatory stage. The dimensionality of the PCA spectrum was also quantified using the participation ratio,

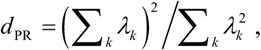

where *λ_k_* are the eigenvalues of the PCA modes. This quantity measures the effective number of modes that contribute substantially to the fluctuations, with smaller values indicating stronger concentration of variance in fewer modes. (A) Sample-averaged *d*_PR_ for all conditions. (B–F) Corresponding curves for GdCl₃, Mid-Small, Near-Head, Near-Foot and Mid-Large tissues, respectively; shaded bands denote bootstrap uncertainty regions. The results are very similar to those obtained with the entropy-based effective dimension *d*_eff_ used in the main text: *d*_PR_ decreases progressively during the preparatory stage in all conditions, the magnitude of the decrease is comparable, and the ordering of the tissue classes is preserved, with GdCl_3_-treated tissues maintaining the highest dimensionality, Mid-Small remaining intermediate, and Mid-Large/ Near-Foot / Near-Head occupying the lower-dimensional regime. Thus, the conclusion that the preparatory stage involves progressive spectral concentration is robust to the specific definition of effective dimensionality.

#### Robustness to contour representation and temporal sampling

To determine whether the decline in effective dimension depends on contour preprocessing or temporal sampling, we repeated the complete analysis while varying the angular resolution of the radial contour representation, the centering convention and the temporal sampling interval. The decline was essentially unchanged when contours were represented by 50, 75 or 100 angular points, or when harmonic centering was replaced by centroid centering (Fig. S6). It also remained significant after subsampling the contours at 5-, 10- and 20-min intervals, although its apparent magnitude decreased as the sampling became coarser. At 20-min sampling, each PCA window contained a median of only eight frames, limiting the number of fluctuation directions that could be resolved. This setting therefore provides a stringent but comparatively low-precision robustness test.

**Fig. S5.**
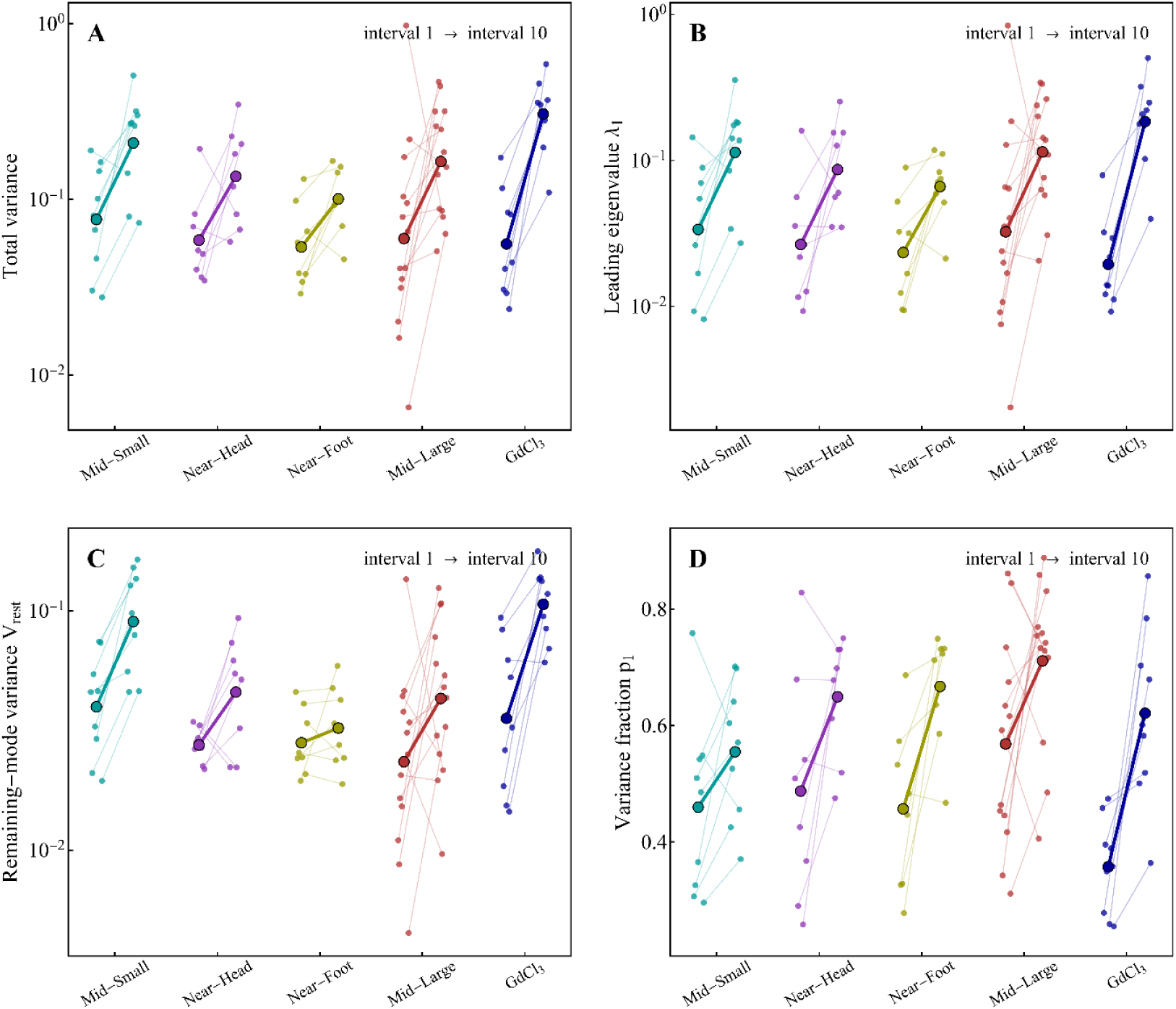
Animal-level changes in fluctuation amplitude and spectral concentration during the preparatory stage. Paired measurements from the first and final equal temporal intervals of the preparatory stage are shown for each tissue. (A) Total fluctuation variance, *V*_tot_ = _∑_*_k_ λ_k_*. (B) Leading PCA eigenvalue, *λ*_1_. (C) Combined variance in the remaining PCA modes, *V*_rest_ = *V*_tot_ − *λ*_1_. (D) Fraction of total variance carried by the leading mode, *P_1_ = λ_1_/V_text_* Small symbols represent individual tissues, and thin lines connect values from the same tissue in intervals 1 and 10. Large outlined symbols and thick lines show condition-level geometric means in A-C and arithmetic means in D. Panels A-C use logarithmic vertical scales, whereas D uses a linear scale. Colors identify the five experimental conditions, as in Fig. 2. In an animal-level analysis in which the five conditions were treated as equally weighted blocks, total variance increased by a factor of 2.82 (blocked-bootstrap 95% confidence interval: 2.19-3.61), *λ*_1_ by a factor of 4.01 (2.91-5.52), and *V*_rest_ by a factor of 1.89 (1.57-2.26), while *p*_1_ increased by 0.175 (0.124-0.225). All four changes were significant in two-sided condition-blocked sign-flip tests after Holm correction (*p* < 10^−4^). Thus, the decline in effective dimension reflects disproportionate amplification of the leading fluctuation direction within a generally expanding fluctuation spectrum.

**Fig. S6.**
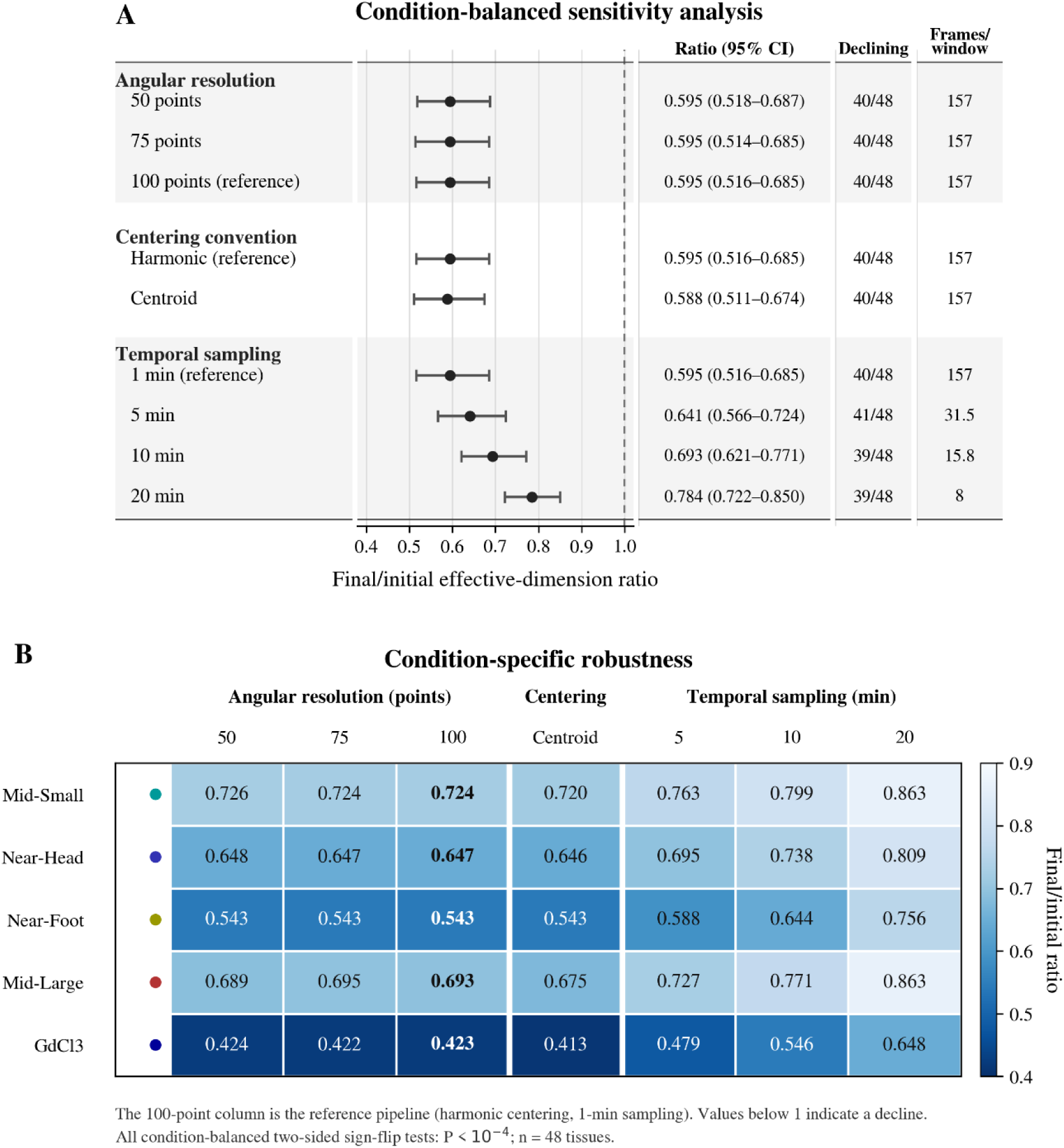
Robustness of the preparatory-stage decline in effective dimension to contour representation and temporal sampling. Effective dimension was recalculated for all 48 tissues after varying the angular resolution of the radial contour representation, the centering convention or the temporal sampling interval. (A) Condition-balanced geometric mean of the ratio between effective dimension in the final and initial normalized temporal intervals. Points and horizontal bars show the estimated ratios and condition-stratified bootstrap 95% confidence intervals. Tissues were resampled separately within each condition, and the five-condition means were assigned equal weight. The dashed line marks a ratio of 1, corresponding to no temporal change. The accompanying columns report the numerical estimate, the number of tissues showing a decline and the median number of frames per PCA window. All condition-balanced two-sided sign-flip tests gave P < 10⁻⁴. (B) Condition-specific geometric mean ratios for the different sensitivity settings. Numbers give the corresponding ratios, with values below 1 indicating a decline. The 100-point column represents the reference analysis, using harmonic centering and 1-min temporal sampling. The decline is insensitive to angular resolution and centering convention and remains evident under all tested temporal subsampling intervals.

### Supplementary Note 2: Temporal correspondence and persistence of PCA modes

The decline in effective dimension during the preparatory stage is accompanied by disproportionate amplification of the leading PCA eigenvalue (Fig. 2G and Fig. S5). This raises the question of whether PCA modes in successive temporal intervals represent related fluctuation directions, or whether each interval selects an unrelated basis. Spectral concentration within individual intervals does not by itself establish temporal persistence of the dominant fluctuation modes. The purpose of the following analysis is to test whether the increasingly concentrated fluctuation spectrum is accompanied by temporal correspondence and progressive stabilization of its leading direction.

The contour representation and its alignment are fixed across the preparatory stage, so PCA eigenvectors obtained in different temporal intervals belong to the same ambient shape space and can be compared directly. Let *e^(r)^_i_* denote the *i*^th^ PCA eigenvector in interval *r*. We quantified correspondence between modes in consecutive intervals using the sign-invariant squared overlap:

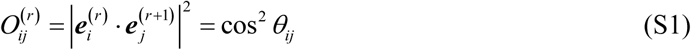

where *θ_ij_* is the angle between the two directions. Thus, *O*_11_ measures persistence of the leading mode, *O*_22_ measures persistence of the second mode, while *O*_12_ and *O*_21_ measure exchange between the first and second modes. Because the sign of a PCA eigenvector is arbitrary, use of the squared scalar product makes these quantities independent of the eigenvector sign convention.

We also calculated the overlap of the complete leading two-dimensional subspaces,

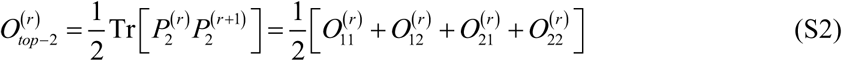

where *P^(r)^_2_* is the projector onto the first two PCA modes in interval *r*. This subspace measure remains well defined when the first two eigenvalues are close and the individual eigenvectors can rotate or exchange order within their common subspace. All overlap measures range from zero, for orthogonal modes or subspaces, to one, for identical modes or subspaces.

We first tested whether neighboring temporal intervals show greater top-two subspace correspondence than intervals that are unrelated in time. This test was performed separately within every tissue. For each tissue, we calculated the top-two overlap between every pair of the 10 preparatory-stage intervals. We then compared the mean overlap of the nine observed consecutive pairs with a null distribution generated by randomly permuting the order of the 10 intervals within that same tissue. No intervals were exchanged between tissues or experimental conditions.

The observed mean adjacent overlap exceeded the mean of the permutation distribution in 46 of the 48 tissues, with positive mean excess in all five experimental conditions. Across tissues, the probability of obtaining this directional consistency under an exact two-sided sign test was *P* 10^−11^. Thus, PCA subspaces in consecutive preparatory-stage intervals are not independent. They retain temporal correspondence despite being estimated separately in each interval.

The permutation test establishes that neighboring intervals retain more similar fluctuation subspaces than intervals that are unrelated in time. However, because it compares the mean overlap of all consecutive pairs with a temporally permuted baseline, it does not test whether the overlap itself increases from early to late intervals. Such an increase could indicate progressive stabilization of the dominant fluctuation direction, but raw overlap cannot be interpreted directly in this way.

Moreover, an apparent increase in overlap can arise even if the underlying fluctuation directions remain completely fixed. As the leading eigenvalue becomes more separated from the remaining spectrum, the corresponding eigenvector can be estimated more precisely from a finite number of frames. Two independent estimates of the same underlying direction will therefore appear more similar at late times than at early times. Because the eigenvalue spectrum becomes increasingly anisotropic during the preparatory stage, this improvement in estimation precision is systematically correlated with time and can mimic progressive stabilization. Testing whether the leading direction genuinely becomes more persistent therefore requires comparing the observed overlap trend with a null model in which the underlying eigenbasis is held fixed while the interval-dependent spectrum, sample size, and temporal correlations are preserved.

We therefore constructed a fixed-basis null model separately for every tissue. First, we pooled all contour fluctuations from the 10 preparatory-stage intervals of that tissue and calculated a single reference PCA basis, {***e****_k_*}. This reference basis represents the time-independent fluctuation directions assumed under the null hypothesis and was used in every simulated interval.

The simulation, nevertheless, preserved the interval-dependent statistical properties that can affect the precision with which these directions are estimated. Let *δx^(r)^_t_* denote the observed centered contour at frame *t* in interval *r*, and let *δx^(r)^_t_* denote the *k*^th^ PCA eigenvector calculated from that interval. The corresponding observed PCA score is defined as

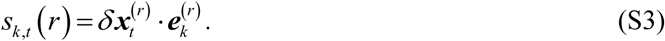

Next, for every interval *r* and mode *k*, we estimated the lag-one autocorrelation as the Pearson correlation between the score at one frame and its value at the following frame:

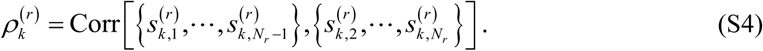

Here, *N_r_* is the number of frames in interval *r*. Thus, *ρ^(r)^_k_* measures the temporal persistence of the experimentally observed amplitude of mode *k* within that interval. To prevent numerically unstable autoregressive simulations, estimated correlations were restricted to the range |ρ^(r)^_k_| ≤ 0.95. If a score contained fewer than four values or had zero variance, its autocorrelation was set to zero.

For every interval, the simulation retained the observed number of frames *N_r_*, the observed PCA eigenvalue *λ^(r)^_k_* and the estimated lag-one autocorrelation *ρ^(r)^_k_* The normalized simulated amplitude of each mode was generated using a first-order autoregressive process:

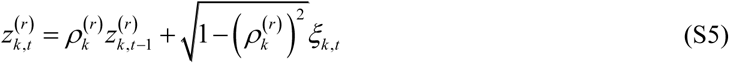

where *ξ_k_* _,*t*_ are independent random variables across modes and frames, each drawn from a normal distribution with zero mean and unit variance. For a fixed value of *ρ^(r)^_k_*, this construction gives *z^(r)^_k,t_* with unit stationary variance and lag-one autocorrelation *ρ^(r)^_k_*. The final simulated value of each mode in one interval was used as the initial value for the next interval, preserving temporal continuity across interval boundaries.

The simulated contour fluctuation was then constructed as

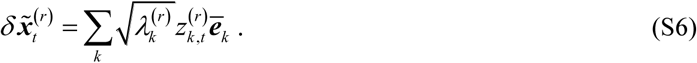

The tilde distinguishes the simulated contour fluctuation from the observed contour fluctuation used to estimate the interval-specific parameters. Thus, the number of observations, fluctuation amplitudes, and temporal correlations varied between intervals as in the experimental data, while the underlying directions ***e****_k_* remained fixed throughout the preparatory stage.

PCA was subsequently recalculated independently in every simulated interval, exactly as for the experimental data. Because each PCA was estimated from a finite and temporally correlated sample, the measured eigenvectors could differ between intervals even though their true underlying directions were constant. One thousand complete simulated trajectories were generated for every tissue. Any increase in their measured overlap therefore represents the increase expected solely from the changing eigenvalue spectrum, finite sampling, and temporal autocorrelation.

As an implementation check, not shown, we repeated the simulation using the observed interval-specific eigenvectors in place of the fixed reference basis. This reconstruction closely reproduced both the magnitude and temporal trajectory of the experimental (*O*_11_), confirming that the simulation and overlap-analysis pipeline recover the measured behavior when the observed basis evolution is supplied. Because the interval-specific eigenvectors are imposed in this reconstruction, it constitutes a sanity check rather than an independent statistical test.

For each overlap quantity, we quantified its temporal trend by fitting a linear model to the nine consecutive interval boundaries:

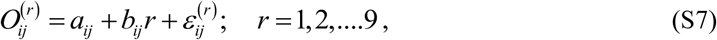

where *a_ij_* and *b_ij_* are fitting parameters which minimize the mean square of the error term *ε^(r)^_k_*. Thus the slope *b_ij_* is the average change in overlap per interval boundary. The observed data provide one slope, *b^obs^_ij_* for each tissue. Each fixed-basis simulation, *m*, similarly provides a null slope, *b^null,m^_ij_*. The mean null slope is the average:

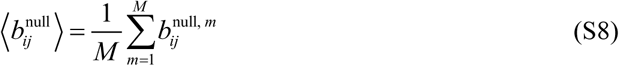

Here, *M* =1,000 is the number of simulated trajectories, each corresponding to an independent realization of the noise *ξ_k_* _,*t*_ in Eq. (S5). We defined the precision-corrected slope as

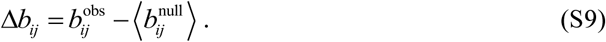

A positive Δ*b_ij_* means that the overlap increases more rapidly in the experimental trajectory than expected from changing eigenvalue spectra, finite sampling, and temporal autocorrelation in a fixed underlying basis.

The observed leading-mode overlap, (*O*_11_), increased during the preparatory stage, but the fixed-basis null also increased substantially, confirming that improving eigenvector precision is an important part of the raw trend (Fig. S7A). Nevertheless, the observed increase exceeded the null expectation. Across the 48 tissues, the animal-weighted mean slope excess was 0.0130, with a condition-stratified bootstrap 95% confidence interval of 0.0028 to 0.0233. When the five experimental conditions were instead assigned equal weight, the mean was 0.0136, with a 95% confidence interval of 0.0038 to 0.0237. The slope excess was positive in 33 of the 48 tissues, corresponding to an exact two-sided sign-test p = 0.013.

The effect was not uniform across conditions (Fig. S7B,C). The clearest positive excess was observed for Near-Head and GdCl₃-treated tissues. Mid-Small and Mid-Large tissues showed positive mean effects with greater animal-to-animal variability, whereas Near-Foot tissues did not show an increase beyond the fixed-basis expectation. Their substantial configuration variability may contribute to this. result by rotating the estimated PCA basis between intervals or making a single eigenbasis an incomplete description of the fluctuations. However, the present analysis does not test this explanation directly

**Fig. S7.**
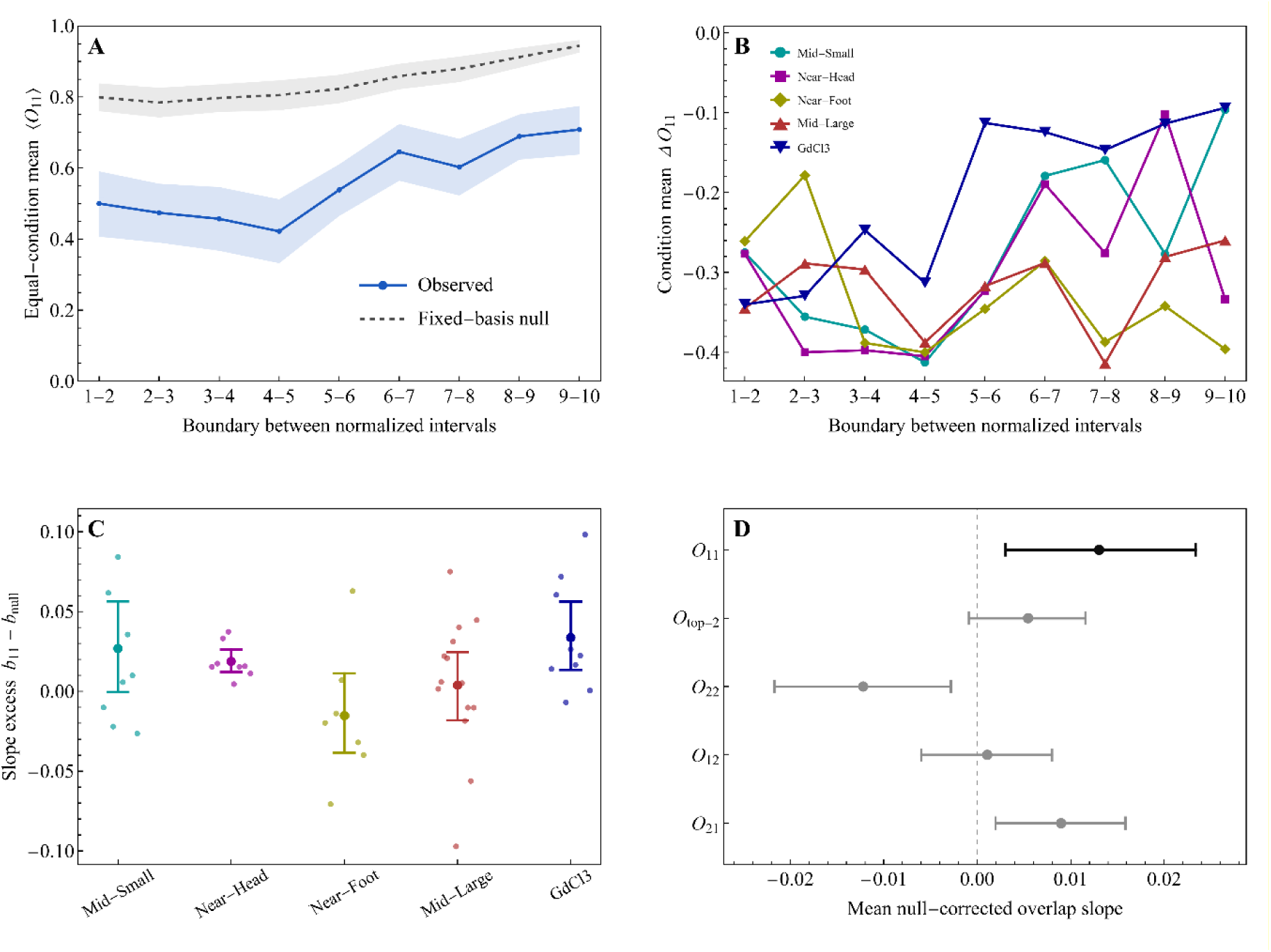
Temporal correspondence and precision-controlled persistence of leading PCA modes during the preparatory stage. PCA modes were calculated independently in 10 normalized temporal intervals, producing nine consecutive-interval overlaps per tissue. The overlap definitions, fixed-basis null model, and statistical procedures are described in Supplementary Note 2. The null model was evaluated using 1,000 simulated trajectories per tissue. (A) Experimental and fixed-basis-null trajectories of the equal-condition mean leading-mode overlap, (*O*_11_). Tissues were first averaged within each condition, and the five condition means were then assigned equal weight. Shaded regions indicate bootstrap 95% confidence intervals. (B) Condition-specific trajectories of the null-corrected leading-mode overlap, Δ*O*_11_, obtained by subtracting the fixed-basis null expectation from the observed *O*_11_ at each interval boundary. (C) Null-corrected slope of *O*_11_ across interval boundaries for individual tissues. Small symbols represent tissues; large symbols and error bars show condition means and bootstrap 95% confidence intervals. (D) Mean null-corrected slopes across tissues for *O*_11_, the top-two projector overlap, *O*_22_, *O*_12_, and *O*_21_. Error bars show condition-stratified bootstrap 95% confidence intervals. *O*_11_ is shown in black and the other overlap measures in gray.

The remaining overlap components did not show the same convergence across statistical summaries (Fig. S7D). In particular, *O*_12_ was centered close to zero, whereas *O*_22_ tended to decrease and *O*_21_ tended to increase. These opposite changes may partly reflect rotation or exchange of the first two estimated modes. The null-corrected increase in the complete top-two projector overlap was small and was not consistently positive across individual tissues. The conclusion is therefore that the leading PCA direction shows modest progressive stabilization beyond the precision baseline, rather than that the complete leading two-dimensional subspace becomes universally more persistent.

These analyses answer two distinct questions. The within-tissue permutation test shows that fluctuation subspaces in neighboring preparatory-stage intervals retain temporal correspondence. The fixed-basis simulation then shows that the leading mode becomes modestly more persistent than expected solely from the evolving eigenvalue spectrum, finite sampling, and temporal autocorrelation. This result supports the interpretation that the observed spectral concentration is accompanied by temporal organization of the leading fluctuation direction.

**Fig. S8.**
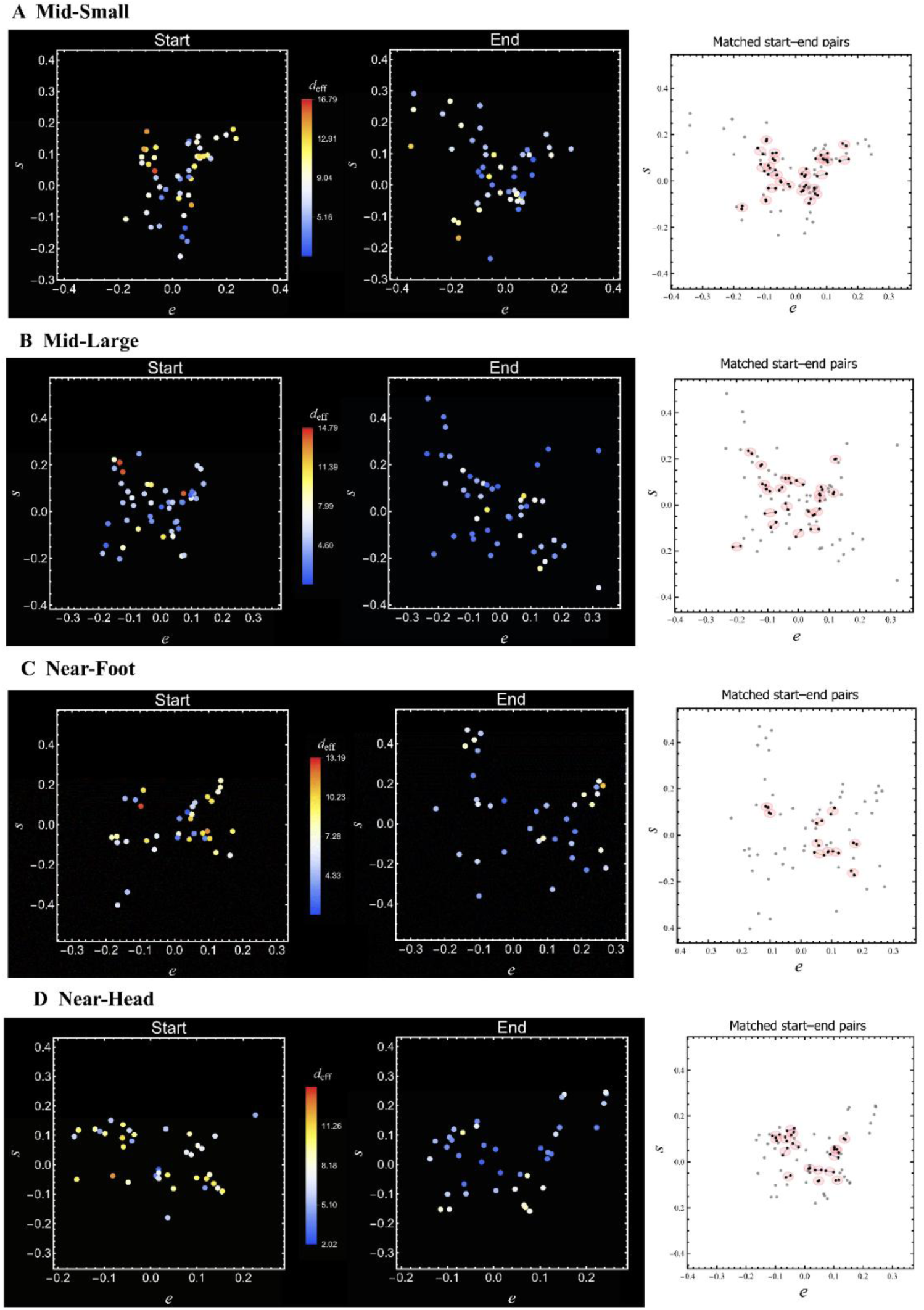
Shape-space maps and matched-shape analysis for the four additional experimental conditions. Rows show Mid-Small, Mid-Large, Near-Foot and Near-Head tissues. The columns show effective-dimension maps at the beginning and end of the early quasi-spherical period, followed by the corresponding initial-final matching map. Analysis and graphical conventions are as in Fig. 4B-D; the pair-radius tolerance was 0.018.

**Table S2.**
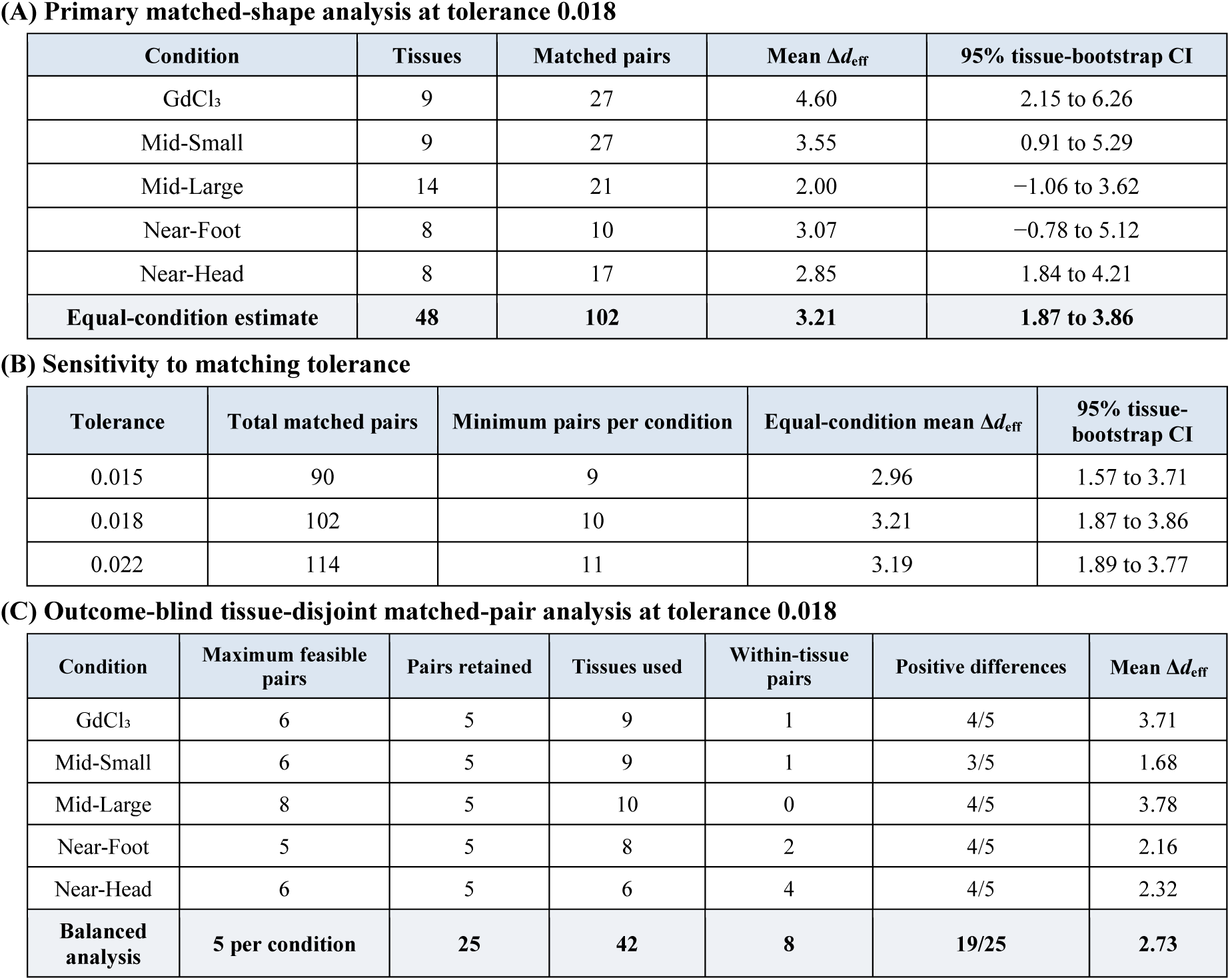
Matched-shape statistics, tolerance sensitivity and independent-pair analysis. (A) Primary matched-shape analysis at tolerance 0.018 (B) Sensitivity to matching tolerance (C) Outcome-blind tissue-disjoint matched-pair analysis at tolerance 0.018

**Notes**. Δd_eff_ = d^start^_eff_ − d^end^_eff_; positive values indicate lower effective dimension in the final period. Matched-pair counts report the number of shape-matched comparisons. Most pairs connect contour sets from different tissues, but each tissue can contribute different contour sets to multiple comparisons; the comparisons were therefore not treated as independent biological replicates. Confidence intervals were obtained by resampling complete tissues within each condition and repeating the matching procedure. The equal-condition estimate assigns equal weight to each of the five condition means.

For the analysis in C, we first identified all pairs containing one contour set from the beginning and one from the end of the preparatory period that satisfied the shape-matching criterion. The pair radius was defined as half the Euclidean separation of the two sets in the (*e*,*s*) shape space. Thus, a tolerance of 0.018 allowed a maximum separation of 0.036. We then required the pairs to be tissue-disjoint: a tissue could contribute to only one pair, either through two sets from the same tissue or through one set matched to a different tissue. This calculation yielded maximum numbers of 6, 6, 8, 5 and 6 independent pairs for GdCl₃, Mid-Small, Mid-Large, Near-Foot and Near-Head, respectively. The largest balanced analysis therefore contained five pairs per condition. For each condition, we selected the feasible set of five pairs having the smallest total separation in shape space. Pair feasibility and selection were determined without using *d*_eff_.

The resulting dataset contained 25 non-overlapping pairs formed from 42 distinct tissues. The five pair differences were averaged within each condition, and the five-condition means were then assigned equal weight. The equal-condition mean Δ*d*_eff_ was 2.73. Its 95% confidence interval, obtained by resampling pairs separately within each condition, was 1.43 to 3.96. An exact two-sided sign-flip test over all 33,554,432 possible assignments of the 25 signs gave *P*<0.001. Nineteen of the 25 differences were positive, giving an exact two-sided sign-test *P*=0.015.

**Fig. S9.**
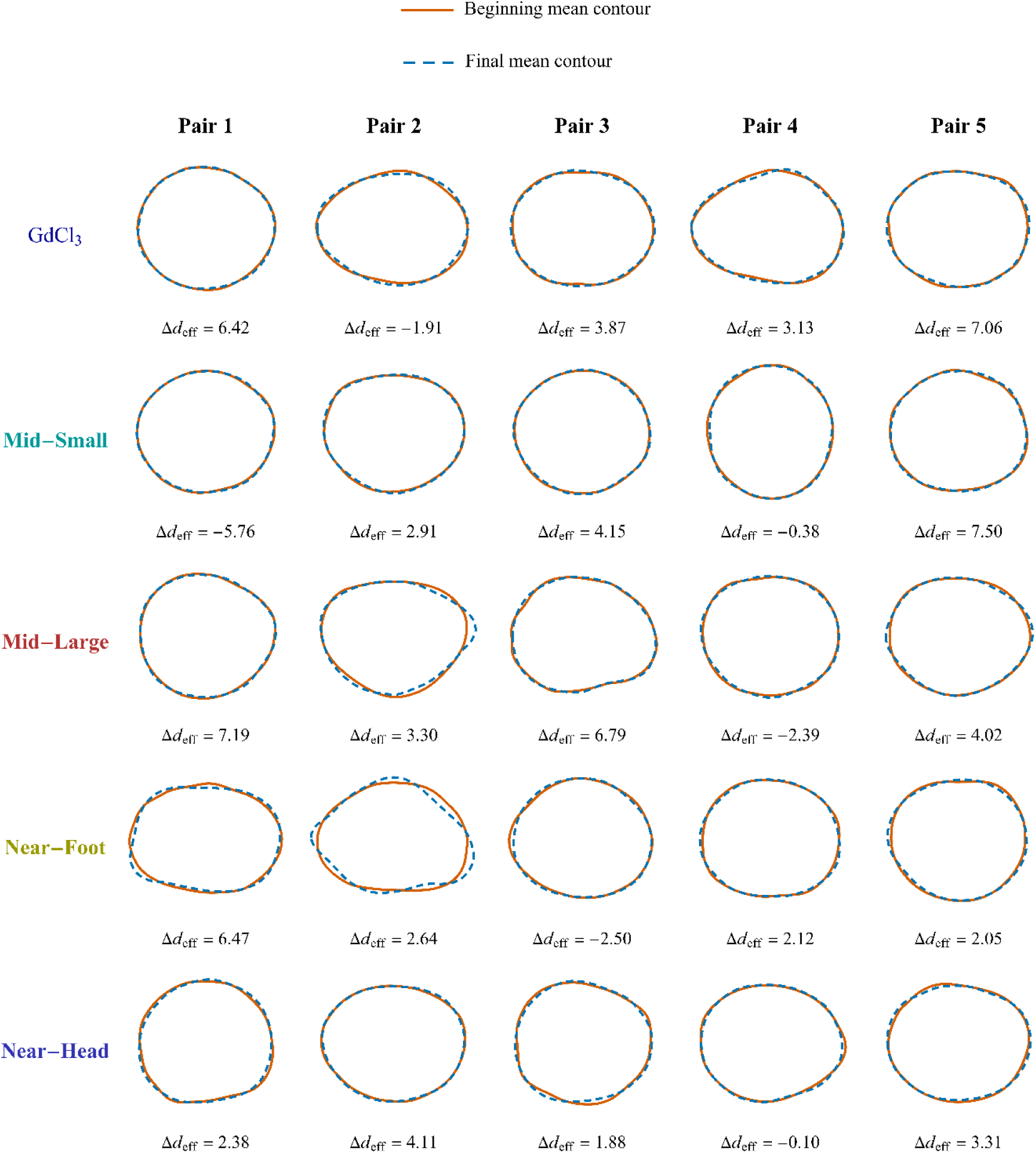
Mean contours of 25 independent shape-matched pairs. Rows correspond to the five experimental conditions and show five tissue-disjoint pairs selected by the outcome-blind shape-matching procedure described in Table S2. Pairs are ordered from left to right by increasing pair radius. The contours were reconstructed from mean radial signatures averaged over approximately 100 consecutive frames and normalized to unit area. Beginning contours are orange and solid; final contours are blue and dashed. Values below the pairs show *Δd_eff_ = d^start^_eff_ − d^end^_eff_*, so that positive values indicate a reduction in effective dimension.

**Fig. S10.**
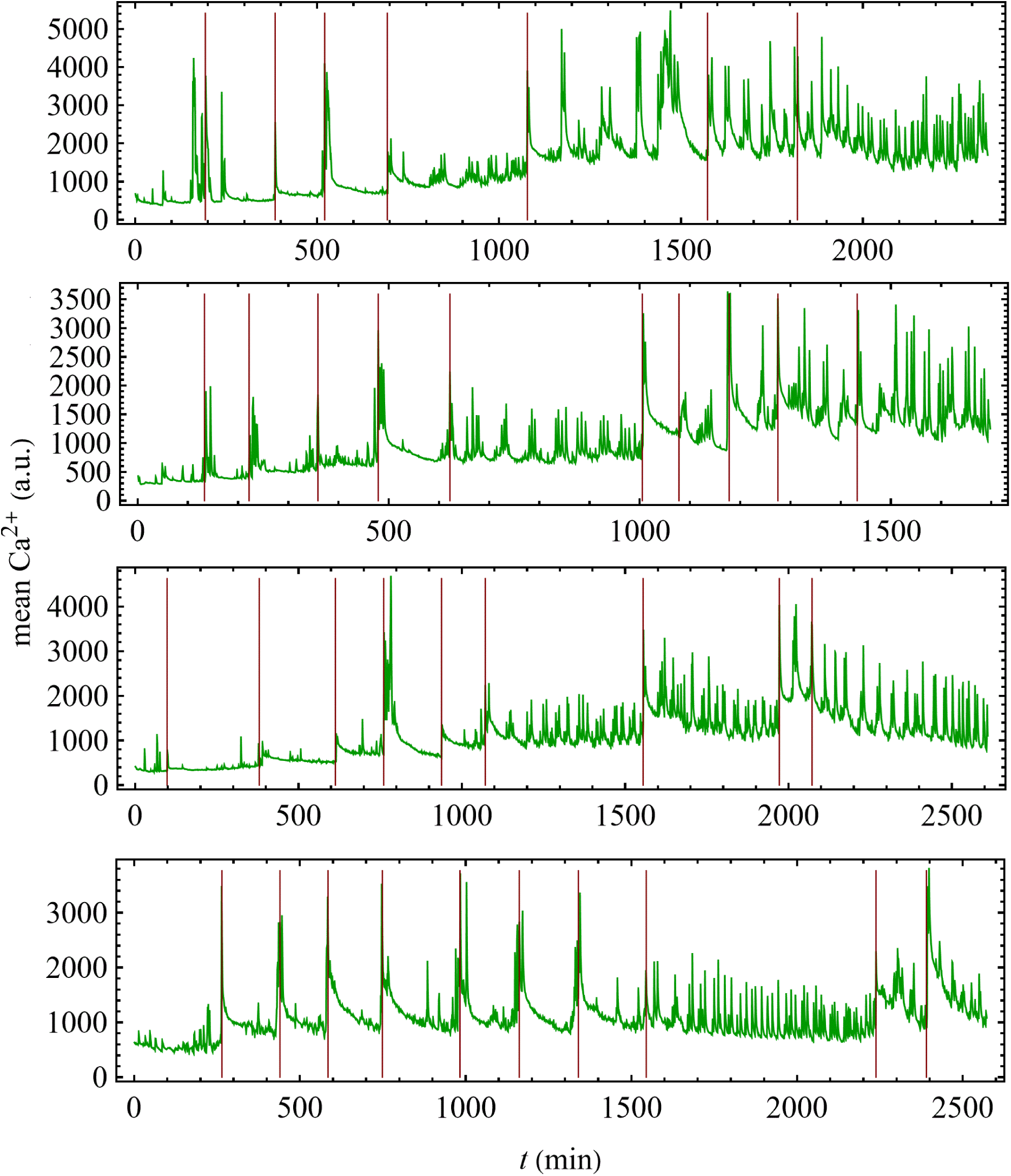
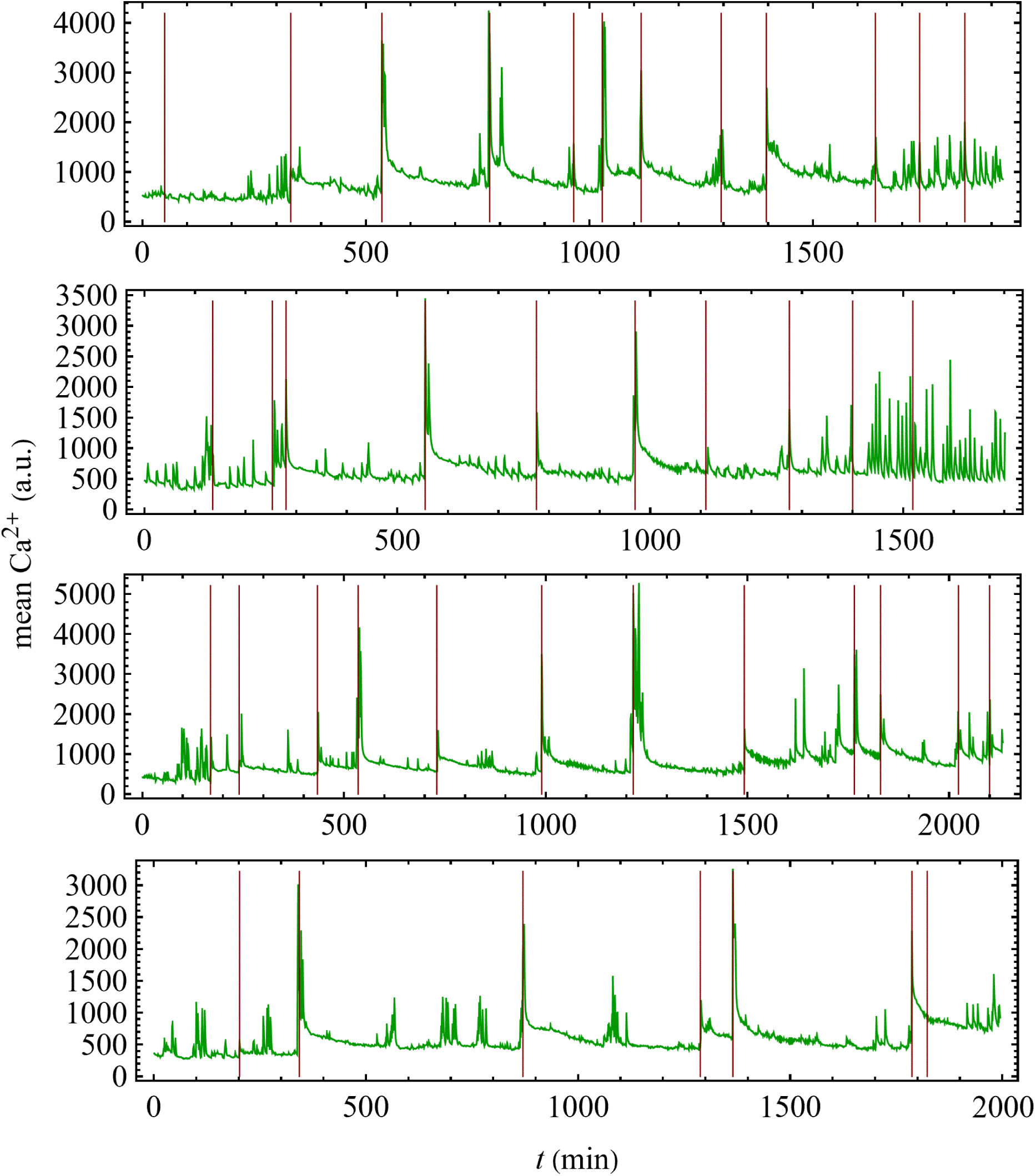
Calcium traces of the GdCl3-treated samples used in the analysis. Shown are the mean Ca^2+^ fluorescence traces for all GdCl_3_-treated tissues analyzed in this work, excluding the representative example displayed in the main text. Green curves show the measured Ca^2+^ signal, and red vertical lines mark the rupture times used to partition the traces into inter-rupture intervals. Together with the example shown in Fig. 5A, these traces illustrate the variability across samples and the common qualitative structure of the calcium dynamics in the GdCl_3_-treated condition, namely rupture-associated calcium elevations superimposed with faster spike-like fluctuations.

### Supplementary Note 3: A model of the calcium activity in GdCl_3_-treated tissues during the preparatory stage

To motivate the interpretation of the experimental calcium traces, we consider a simple analytic model for the calcium activity in GdCl_3_-treated tissues. The model reflects the electrically excitable nature of *Hydra* tissue leading to Ca^2+^ excitations, akin to the characteristics of a smooth muscle[4-7]. The nature of the Ca^2+^ dynamics is determined by a few ingredients: the existence of an activation threshold and a wide range of the Ca^2+^ relaxation times, reflecting the multiple intracellular processes governing calcium dynamics[7-9]. In this model, the measured Ca^2+^ signal is represented as the sum of two contributions: a rupture-associated slowly-relaxing background component and a faster spike-like component. The background component changes through discrete upward jumps at rupture events and relaxes more gradually between them, thereby defining the interval-scale calcium state of the tissue. Superimposed on this evolving background is a faster excitable component that gives rise to the spike-like activity. A representative simulation of the model is shown in Fig. S11. The purpose of this toy model is not to reproduce quantitatively in every detail the experimental traces, but rather to capture the qualitative separation between rupture-associated background dynamics and fast spike-like events, and to provide a framework for interpreting the relation between Ca^2+^ level, excitability, and morphology. In particular, the model is intended to describe the GdCl_3_-treated condition, in which the rupture-associated calcium background appears to relax unusually slowly compared with untreated tissues.

**Fig. S11.**
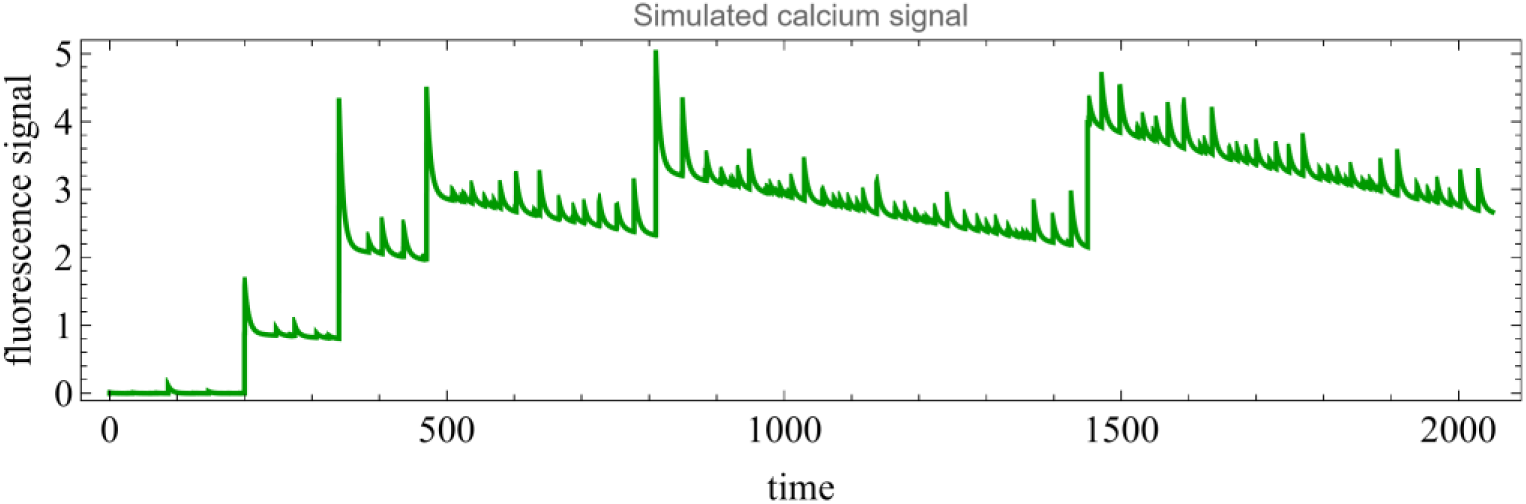
Representative simulated calcium trace from the analytic model. The simulated signal consists of a rupture-associated background component that undergoes abrupt upward jumps at rupture events and relaxes gradually between them, together with a faster spike-like component superimposed on this background. The trace illustrates the qualitative separation between interval scale calcium modulation and local spike-like events used to interpret the experimental data. For the simulation shown here, the model parameters were τ_R_ =1500, τ_exc_=6, τ_H_=10, α=2, γ=10, σ=3, B=14, β=7, x_0_=2.5. The kick amplitudes were taken to be equal, *A_k_* = 1.5(*r_k_* + 0.5), where *r_k_* are independent random variables uniformly distributed between zero and one. The comparatively large value of τ_R_ was chosen to reflect the slow relaxation of the rupture-associated calcium background observed in GdCl_3_-treated tissues.

In its simplest form, the model consists of a rupture-associated background component, *R* (*t*), which undergoes discrete upward kicks at rupture events and then relaxes slowly between them. The spike-like contribution is represented by a single excitable variable, *c* (*t*), which is coupled to *R* (*t*) and opposed by a recovery variable *h* (*t*). The measured fluorescence signal is assumed to be proportional to the sum of these two components, *C* (*t*) = *R* (*t*) + *c* (*t*).

The rupture-associated component evolves according to

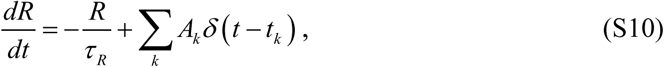

where *τ _R_* is the slow relaxation time, and *A_k_* is the amplitude of the calcium kick produced by rupture event *k* occurring at time *t_k_*. In the present context, *τ _R_* is taken to be long, consistent with the observation that in GdCl_3_-treated tissues the rupture-associated calcium background decays slowly between successive rupture events.

The spike-like component satisfies

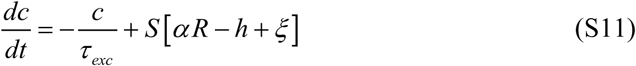

where *τ _exc_*□*τ _R_* is a much shorter relaxation time, *α* controls the coupling of the excitable component to the background mode, and *ξ* (*t*) is a stochastic forcing term taken to be Gaussian white noise with zero mean and variance set by *ξ* (*t*)*ξ* (*t*′) = *σ* ^2^*δ* (*t* − *t*′).

The nonlinear activation function is chosen to be a sigmoid,

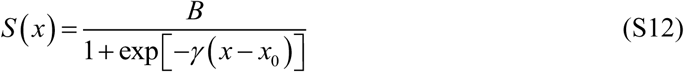

where *B* is the maximal amplitude, *x*_0_ sets the activation threshold, and *γ* determines the steepness of the transition.

Finally, the recovery variable obeys

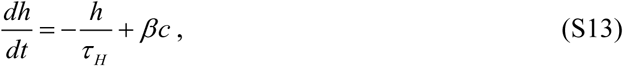

where *β* determines the strength of the recovery term, and *τ _H_* its characteristic time scale. Together, these equations provide a minimal analytic description of the calcium signal as the sum of a rupture-associated background and a spike-like excitable component.

Within this model, rupture events do not directly generate the fast spike-like fluctuations themselves. Rather, each rupture event produces an upward kick in the background component *R*(*t*). Because the relaxation of *R*(*t*) is slow compared with the interval between successive ruptures, these kicks accumulate over time and progressively elevate the background calcium level. As a result, the effective drive to the excitable system gradually increases, bringing it closer to threshold. Spike-like activity, therefore, tends to appear only later, once the accumulated background has become sufficiently large so that stochastic fluctuations can trigger pronounced excursions of *c*(*t*). In the context of GdCl_3_-treated tissues, this accumulation picture is particularly natural: if GdCl_3_ slows the relaxation of the rupture-associated calcium background, then successive rupture-induced kicks will be less effectively dissipated, allowing the background level to build up over time and thereby promoting the delayed emergence of spike-like calcium events. In this way, the model captures the central qualitative feature that the rupture-associated background controls the excitability state of the tissue and thereby enables the emergence of spike-like calcium events. Note that although GdCl_3_ blocks external Ca^2+^ channels, in particular stretch-activated channels, the cytoplasmic Ca^2+^ can increase due to enhanced fluxes from internal reservoirs. Thus, the GdCl_3_ does not necessarily mean a reduction in overall Ca^2+^ activity but rather slowing-down or decrease in the calcium response to mechanical deformations of the tissue (see [1] for discussion of this point).

This picture motivates the decomposition used in the experimental analysis. The purpose of the decomposition is not to claim that the measured Ca^2+^ traces follow the model in a literal quantitative sense, but rather to extract from the data the two qualitative ingredients suggested by the model: a rupture-associated background component that evolves on the interval scale through successive rupture events, and a faster residual spike-like component superimposed on that background.

## References

1. Gilbert, S.F., and Barresi, M.J.F. (2016). Developmental Biology, 11 Edition (Oxford University Press).

2. Waddington, C.H. (1942). Canalization of development and the inheritance of acquired characters. Nature 150, 563–565.

3. Waddington, C.H. (1957). The Strategy Of The Genes (Allen and Unwin).

4. Mammoto, T., Mammoto, A., and Ingber, D.E. (2013). Mechanobiology and Developmental Control. Annual Review of Cell and Developmental Biology 29, 27–61.

5. Braun, E., and Keren, K. (2018). Hydra Regeneration: Closing the Loop with Mechanical Processes in Morphogenesis. BioEssays 40, 1700204.

6. Lecuit, T., Lenne, P.-F., and Munro, E. (2011). Force generation, transmission, and integration during cell and tissue morphogenesis. Annual review of cell and developmental biology 27, 157–184.

7. Livshits, A., Shani-Zerbib, L., Maroudas-Sacks, Y., Braun, E., and Keren, K. (2017). Structural inheritance of the actin cytoskeletal organization determines the body axis in regenerating hydra. Cell Reports 18, 1410–1421.

8. Miller, C.J., and Davidson, L.A. (2013). The interplay between cell signaling and mechanics in developmental processes. Nature Rev. Genetics 14, 733–744.

9. Howard, J., Grill, S.W., and Bois, J.S. (2011). Turing’s next step: the mechanochemical basis of morphogenesis. Nature Review Molecular Cell Biology 12, 392–398.

10. Braun, E., and Ori, H. (2019). Electric-Induced Reversal of Morphogenesis in Hydra. Biophysical Journal 117, 1514–1523.

11. Agam, O., and Braun, E. (2023). Universal Calcium fluctuations in Hydra morphogenesis. Phys. Biol. 20, 066002

12. Levin, M. (2021). Bioelectric signaling: Reprogrammable circuits underlying embryogenesis, regeneration, and cancer. Cell 184, 1971–1989.

13. Agam, O., and Braun, E. (2025). Stochastic morphological swings in Hydra regeneration: A manifestation of noisy canalized morphogenesis. Proceedings of the National Academy of Sciences 122, e2415736121.

14. Agam, O., and Braun, E. (2023). Hydra morphogenesis as phase-transition dynamics. EPL 143, 27001.

15. Maroudas-Sacks, Y., and Keren, K. (2021). Mechanical Patterning in Animal Morphogenesis. Annu Rev Cell Dev Biol. 37,: 469–493.

16. Agam, O., and Braun, E. (2026). The Dual Nature of Body-Axis Formation in Hydra Regeneration: Polarity-Morphology Concurrency. Biophys J. 125, 3328–3340.

17. Galliot, B. (2012). Hydra, a fruitful model system for 270 years. Int. J. Dev. Biol. 56, 411–423.

18. Galliot, B. (2013). Regeneration in Hydra; Encyclopedia of Life Sciences (John Wiley & Sons Ltd).

19. Bode, H.R. (2009). Axial Patterning in Hydra. Cold Spring Harbor Perspectives in Biology a000463, 1.

20. Maroudas-Sacks, Y., Garion, L., Shani-Zerbib, L., Livshits, A., Braun, E., and Keren, K. (2021). Topological defects in the nematic order of actin fibres as organization centres of Hydra morphogenesis. Nature Physics 17, 251–259.

21. Cummings, S.G., and Bode, H.R. (1984). Head regeneration and polarity reversal in Hydra attenuata can occur in the absence of DNA synthesis. Wilhelm Roux’s archives of developmental biology 194, 79–86.

22. Gierer, A., Berking, S., Bode, H., David, C.N., Flick, K., Hansmann, G., Schaller, C.H., and Trenkner, E. (1972). Regeneration of hydra from reaggregated cells. Nature/New Biology, 98– 101.

23. Park, H.D., Ortmeyer, A.B., and Blankenbaker, D.P. (1970). Cell division during regeneration in Hydra.

24. Kucken, M., Soriano, J., Pullarkat, P.A., Ott, A., and Nicola, E.M. (2008). An osmoregulatory basis for shape oscillations in regenerating Hydra. Biophys J 95, 978–985.

25. Wang, R., Bialas, A.L., Goel, T., and Collins, E.S. (2023). Mechano-Chemical Coupling in Hydra Regeneration and Patterning. Integr Comp Biol. 63, 1422–1441.

26. Ferenc, J., Papasaikas, P., Ferralli, J., Nakamura, Y., Smallwood, S., and Tsiairis, C.D. (2021). Mechanical oscillations orchestrate axial patterning through Wnt activation in Hydra. Science Advances 7, eabj6897.

27. Shani-Zerbib, L., Garion, L., Maroudas-Sacks, Y., Braun, E., and Keren, K. (2022). Canalized Morphogenesis Driven by Inherited Tissue Asymmetries in Hydra Regeneration. Genes 13, 360.

28. Maroudas-Sacks, Y., Suganthan, S., Garion, L., Ascoli-Abbina, Y., Westfried, A., Dori, N., Pasvinter, I., PopoviÄ‡, M., and Keren, K. (2025). Mechanical strain focusing at topological defect sites in regenerating Hydra. Development. 15, 152.

29. Agam, O., and Braun, E. (2026). Rupture-Repair Cycles in Regenerating Hydra Tissues. bioRxiv, 2026.2003.2002.708976. doi: 10.64898/2026.03.02.708976.

30. Agam, O., and Braun, E. (2024). Fluctuation-driven morphological patterning: An unconventional approach to morphogenesis. Physical Review Research 6, 043027.

31. Del Giudice, M. (2021). Effective Dimensionality: A Tutorial. Multivariate Behavioral Research 56, 527–542.

32. Lefebvre, M.F., Claussen, N.H., Mitchell, N.P., Gustafson, H.J., and Streichan, S.J. (2023). Geometric control of myosin II orientation during axis elongation. eLife 12, e78787. 10.7554/eLife.78787.

33. Mitchell, N.P., Cislo, D.J., Shankar, S., Lin, Y., Shraiman, B.I., and Streichan, S.J. (2022). Visceral organ morphogenesis via calcium-patterned muscle constrictions. eLife 11, e77355. 10.7554/eLife.77355.

34. Streichan, S.J., Lefebvre, M.F., Noll, N., Wieschaus, E.F., and Shraiman, B.I. (2018). Global morphogenetic flow is accurately predicted by the spatial distribution of myosin motors. eLife 7, e27454.

35. Szymanski, J., and Yuste, R. (2019). Mapping the Whole-Body Muscle Activity of Hydra vulgaris. Current Biology 29, 1807–1817.

36. Coste, B., Mathur, J., Schmidt, M., Earley, T.J., Ranade, S., Petrus, M.J., Dubin, A.E., and Patapoutian, A. (2010). Piezo1 and Piezo2 are essential components of distinct mechanically activated cation channels. Science 330, 7–12.

37. Ermakov, Y.A., Kamaraju, K., Sengupta, K., and Sukharev, S. (2010). Gadolinium ions block mechanosensitive channels by altering the packing and lateral pressure of anionic lipids. Biophys J. 98, 1018–1027.

38. Alzugaray, M.E., Gavazzi, M.V., Griffo, L., and Ronderos, J.R. (2025). Piezo proteins, mechano reception and behaviour in Hydra. Scientific Reports 15, 6440.

39. Salleo, A., La Spada, G., and Barbera, R. (1994). Gadolinium is a powerful blocker of the activation of nematocytes of Pelagia noctiluca. J Exp Biol. 187, 201–206.

40. Skokan, T.D., Hobmayer, B., McKinley, K.L., and Vale, R.D. (2024). Mechanical stretch regulates macropinocytosis in Hydra vulgaris. Mol Biol Cell. 35, br9.

41. Seroussi, I., Veikherman, D., Ofer, N., Yehudai-Resheff, S., and Keren, K. (2012). Segmentation and tracking of live cells in phase-contrast images using directional gradient vector flow for snakes. Journal of Microscopy 247, 137–146.

42. Pincus, Z., and Theriot, J.A. (2007). Comparison of quantitative methods for cell-shape analysis. Journal of Microscopy 227, 140–156.

## References

1. Agam, O., and Braun, E. (2026). Rupture-Repair Cycles in Regenerating Hydra Tissues. bioRxiv, 2026.2003.2002.708976.

2. Kucken, M., Soriano, J., Pullarkat, P.A., Ott, A., and Nicola, E.M. (2008). An osmoregulatory basis for shape oscillations in regenerating Hydra. Biophys J 95, 978–985.

3. Livshits, A., Shani-Zerbib, L., Maroudas-Sacks, Y., Braun, E., and Keren, K. (2017). Structural inheritance of the actin cytoskeletal organization determines the body axis in regenerating hydra. Cell Reports 18, 1410–1421.

4. Agam, O., and Braun, E. (2023). Universal Calcium fluctuations in Hydra morphogenesis. Phys. Biol. 20, 066002

5. Braun, E., and Ori, H. (2019). Electric-Induced Reversal of Morphogenesis in Hydra. Biophysical Journal 117, 1514–1523.

6. Szymanski, J., and Yuste, R. (2019). Mapping the Whole-Body Muscle Activity of Hydra vulgaris. Current Biology 29, 1807–1817.

7. Wang, H., Swore, J., Sharma, S., Szymanski, J., Yuste, R., Daniel, T., Regnier, M., Bosma, M., and Fairhall, A.L. (2023). A complete biomechanical model of Hydra contractile behaviors, from neural drive to muscle to movement. PNAS 120, e2210439120.

8. Berridge, M.J. (2008). Smooth muscle cell calcium activation mechanisms. The Journal of Physiology 586, 5047–5061.

9. Kuo, I.Y., and Ehrlich, B.E. (2015). Signaling in muscle contraction. Cold Spring Harbor perspectives in biology 7, a006023–a006023.

